# Engineering Resilient Gene Drives Towards Sustainable Malaria Control: Predicting, Testing and Overcoming Target Site Resistance

**DOI:** 10.1101/2024.10.21.618489

**Authors:** Ioanna Morianou, Lee Phillimore, Bhavin S. Khatri, Louise Marston, Matthew Gribble, Austin Burt, Federica Bernardini, Andrew M. Hammond, Tony Nolan, Andrea Crisanti

## Abstract

CRISPR-based gene drives are selfish genetic elements with the potential to spread through entire insect populations for sustainable vector control. Gene drives designed to disrupt the reproductive capacity of females can suppress laboratory populations of the malaria mosquito. However, any suppressive intervention will inevitably exert an evolutionary pressure for resistance. Here, we present a pipeline for the accelerated discovery, engineering, and testing of both natural and drive-induced variants that could reverse gene drive spread. We applied our method to stress-test a highly effective gene drive that has evaded resistance in all laboratory-contained releases to date, known as Ag(QFS)1. We showed that previously undetected resistant alleles can arise at low frequency, and discovered novel, partially resistant alleles that can perturb drive-invasion dynamics. We then engineered next-generation gene drives that can actively remove resistant alleles by targeting several highly conserved and non-overlapping sites in the female-specific exon of the *doublesex* gene. Our models predict that such gene drive designs could suppress large, natural populations of the malaria mosquito in the field.

## INTRODUCTION

Synthetic gene drives can be used to spread desirable traits through natural pest or disease vector populations for sustainable genetic control. The most advanced gene drives to date are based on CRISPR components, and they bias their inheritance by converting the germline of a gene drive heterozygote to homozygosity. Briefly, they comprise genes encoding a Cas9 and gRNA inserted within their own recognition site in a chromosome, so that their expression promotes cleavage of the wild-type chromosome followed by homology-directed repair (HDR) using the gene drive chromosome as a template – a process called homing (**Supp. Figure 1A**). These have been adapted for use in fruitflies^1^, mice^2^ and mosquitoes^3–6^, and have been shown to spread autonomously from low initial frequency^7–9^.

Gene drives can be programmed to reduce the breeding potential of a species by disrupting target genes essential for female reproduction^4,10^. This strategy of population suppression shows great promise for control of the African malaria mosquito, *Anopheles gambiae*^7–9^.

A major technical obstacle in developing these systems is target site resistance, whereby naturally occurring genetic variation or drive-induced mutations, introduced through error-prone repair of the cut site (**Supp. Figure 1A**), prevent further cleavage by the gene drive^11–13^. The extent to which these alleles can limit gene drive spread depends on their initial frequency in the natural target population, how often they are generated, and whether they confer a fitness cost^14–16^. Consequently, the nature, creation and selection of resistance has been the object of intense investigation.

Resistant variants are distinguished into two categories depending on whether they restore a functional copy of the target gene (**R1**) or not (**R2**)^11,17^. **R1** alleles block homing, conferring no fitness cost, and can therefore be subject to strong positive selection in the face of a gene drive, reversing its spread^11,13,17^. Conversely, **R2** alleles block homing but confer strong fitness costs, such as complete sterility or inviability in homozygosity^17,18^. Depending on their frequency, and the fitness of drive-carrying individuals, **R2** alleles can perturb gene drive invasion dynamics, and in some cases form a stable equilibrium that could reduce the magnitude of population suppression^19^. However, their effect on gene drive spread is substantially less severe than that of functional resistance (**R1**) and, in most cases, **R2** alleles are not expected to be a significant barrier to gene drive efficacy^11,17^.

The primary source of novel resistance is error-prone end-joining (EJ), which can occur in the fraction of cleaved chromosomes that do not repair by HDR (**Supp. Figure 1A**). In contrast to the mosquito germline where HDR is dominant, cleavage events in the embryo are strongly biased towards EJ. Thus, a major source of resistance is maternal deposition of nuclease into the embryo^3,11,12,17,20^. Restricting Cas9 expression to the early germline has proven an effective mitigation strategy, reducing the total production of resistant alleles^17,20,21^. Resistance can be further mitigated by targeting functionally constrained sites that permit little or no sequence diversity, biasing EJ towards the production of **R2** over **R1** alleles and reducing the likelihood of naturally occurring resistant variants^22,23^.

We previously developed a population suppression drive, targeting the *doublesex (dsx)* gene^7^ in *An. gambiae,* that was designed to mitigate resistance by combining the above strategies: i.e. germline-restricted Cas9 expression with a functionally constrained and highly conserved target site (**Supp. Figure 1B**). This strain, known as Ag(QFS)1^9^, carries a drive allele that disrupts the female-specific exon of *dsx* (ex5), impairing female development when homozygous, leading to a sterile intersex phenotype^7^. The gene drive was shown to spread rapidly in laboratory-containment, in both small cages, and large cages imitating natural conditions, causing complete population suppression without selection for resistance^7,9^. However, given that natural mosquito populations are far larger than those of the lab (>10^6^ vs ∼10^3^), even very rare **R1** alleles may have the opportunity to be selected in the field (**Supp. Figure 1C**). Thus, efforts to predict, pre-empt, and mitigate resistance prior to a first gene drive release are warranted^16,22^.

To better understand the potential for resistance at the highly conserved target site in *dsx*, we developed a pipeline to discover, engineer and test both natural and drive-induced variants. Our approach simulates gene drive-induced EJ repair at scale, revealing rare resistant alleles and their relative frequencies. We show experimentally that there is evolutionary space for functional **R1** alleles to arise against the Ag(QFS)1 gene drive, and that these can come under positive selection in laboratory populations.

We demonstrate that resistant alleles can be actively removed by multiplexing the gene drive so that it recognises additional non-overlapping sites in *dsx* ^10,14,21,24^ (**Supp. Figure 2**). Our models predict that this approach could allow the suppression of large, natural populations in the field.

Our data and workflow serve as a paradigm for designing and testing the robustness of population suppression gene drives prior to release in the field.

## RESULTS

To determine the evolutionary space for resistance to evolve at the region of *dsx* targeted by Ag(QFS)1^7^, we focused on two sources of variation: standing variation detected in the field; and drive-induced variation, created by end-joining repair following Cas9-induced cleavage. Our approach was to then recreate and test these variants for their ability to block gene drive activity.

### Standing variation among natural mosquito populations may harbour resistant alleles

In *An. gambiae* the best performing population suppression gene drive strain to date targets a site (T1) at the boundary between intron 4 and exon 5 of the *dsxF* isoform, chosen for its high level of sequence conservation across species in the *Anopheles* genus^25^ (**Supp. Figure 3A**) and among wild caught mosquitoes sequenced as part of the first release of the *Anopheles gambiae* 1000 genomes (Ag1000G) project^26^, where only a single SNP was reported (GèA, 2R:48714641). *In vitro* experiments suggested this SNP may still be susceptible to cleavage by the Cas9 present in the gene drive^7,27^. Nonetheless, anticipating the need for second-generation gene drives targeting multiple sites, we expanded our search to 2 additional, non-overlapping potential target sites on the female-specific exon 5 and examined standing variation at all 3 sites, using phase 3 Ag1000G data that covered an extended sample size of >2,700 African malaria mosquitoes, comprising samples of *An. gambiae*, *An. coluzii* and *An. arabiensis* collected from 19 countries (**Supp. Figure 3A**)^26^.

Across the three target sites, T1-T3, 7 SNPs were identified, 6 of which were found in only 2 or fewer individuals (**Supp. Figure 3B**). The only SNP at an appreciable frequency was the original G→A SNP found in T1,at 1.3% overall allelic frequency across all collection sites (**Supp. Figure 3B; Supp. Figure 4**), but reaching as high as 25.9% in Angola, and appears to be in Hardy-Weinberg equilibrium, meaning that it is likely under no or minimal negative selection. Since this mutation is likely to preserve the function of the *dsx* gene it was prioritised for resistance testing.

### Drive-induced EJ can be simulated in a high-throughput mutagenesis screen to identify putatively resistant alleles at *dsx*

End-joining repair of chromosomes cleaved by the gene drive nuclease can be a more important source of resistant mutations. Investigating such drive-induced resistance can be challenging, particularly in a strain like Ag(QFS)1, because high rates of HDR in the germline leave very few cleavage events that are resolved by end-joining (EJ). Thus, insects carrying putatively resistant alleles are extremely rare. Using rates of homing and EJ measured for Ag(QFS)1 (**Supp. Data File 1**)^7^, we estimate that 1.75% of the progeny of drive heterozygotes will carry an EJ mutation. Thus, fewer than 200 EJ events would be expected amongst 10,000 progeny of drive heterozygotes.

Previous research has revealed that, in the embryo, most cleavage events are repaired by end-joining. Consequently, maternally deposited nuclease can contribute to high rates of gene-drive resistant mutations being generated in the germline^3,4,11,17,21,28^; in some cases, over 70% of non-drive alleles in the offspring can carry an EJ mutation^17^.

We set out to simulate and enhance drive-induced EJ mutations at the target of Ag(QFS)1, and subsequently isolate the fraction of *dsx* alleles that may be resistant to gene drive activity. First, we created a new strain with the Ag(QFS)1 allele (comprising Cas9 and gRNA targeted to *dsx*) integrated away from its target site (“*dsx*-mutator” strain) so as to remove the possibility for homing. Next, to enhance EJ through maternal deposition, we crossed males of the *dsx*-mutator strain to females expressing Cas9 under the *vasa2* promoter (“maternal Cas9” strain^3,17,21,28^). This approach generated high rates of mutagenesis in the F2, evidenced by close to 100% frequency of mosaic intersex in F2 females carrying the *dsx*-mutator allele and deposited Cas9 (**Figure 1A, Supp. Figure 5-6**). However, high rates of intersex could be caused by just one or several EJ events in early embryogenesis. Thus, to estimate of the minimum number of cleavage events occurring in the germline, we examined the progeny of 8 F2 males for the presence of distinct EJ alleles. We found that 96% of alleles were mutated, and that mutations were clustered with at least 1-4 different alleles in each clutch (10 F3 individuals sequenced / clutch). These results are consistent with deposited nuclease activity occurring in the small number of stem-cell progenitors present during early embryonic development, prior to expansion of the germline (**Supp. Figure 7**)^29^.

**Figure 1.**
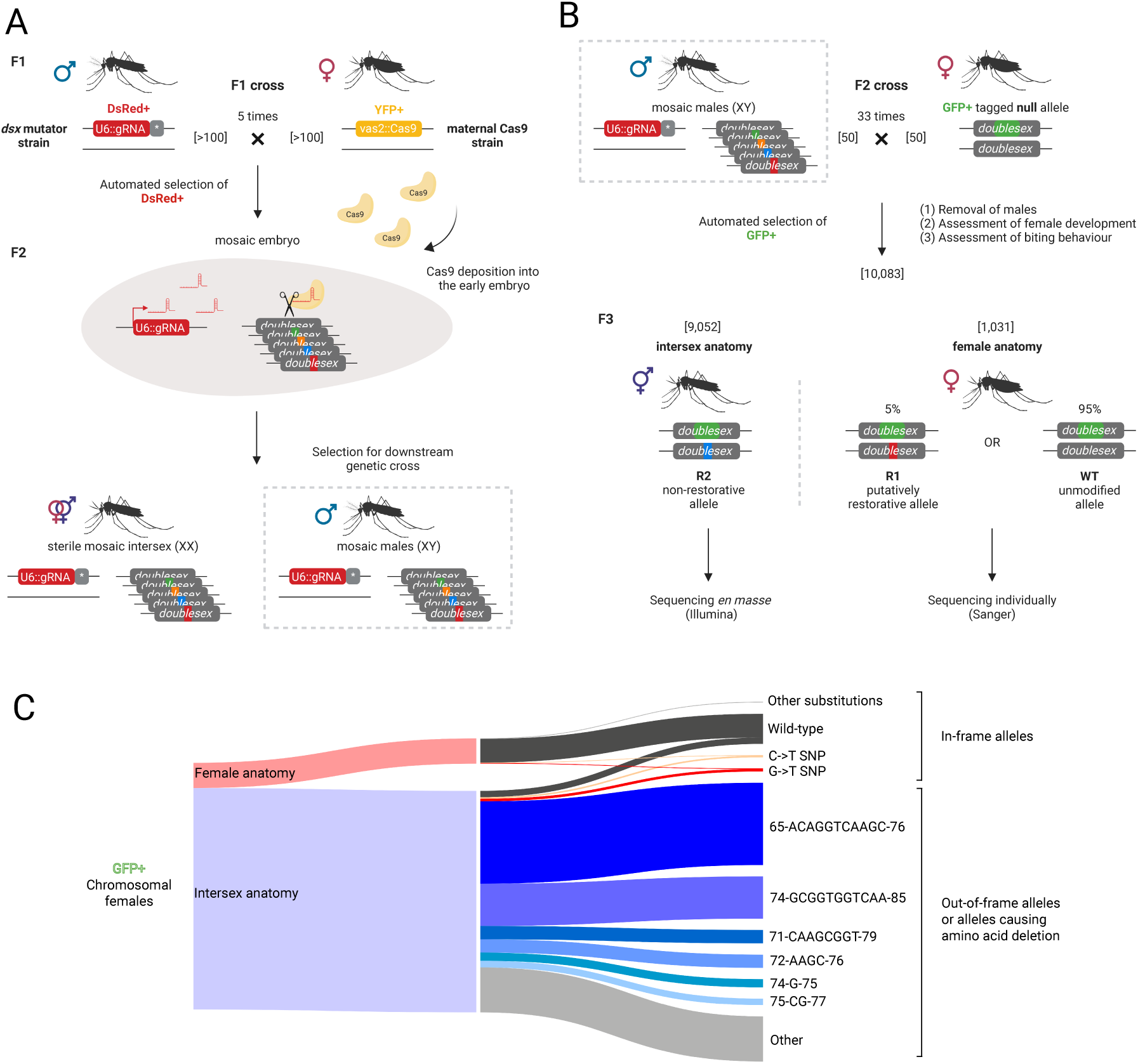
A high throughput mutagenesis screen used to generate and assess Cas9-induced mutations revealed two putative resistant alleles at the target site of Ag(QFS)1. **(A)** A minimum of 100 males of a *dsx* mutator strain (DsRed+) expressing a ubiquitous gRNA against the Ag(QFS)1 target site and a linked *zpg*-Cas9 (grey box), were crossed to a minimum of 100 females expressing *vas2::Cas9* (YFP+) (F1 cross, repeated 5 times). DsRed+YFP-offspring were selected using the COPAS larval sorter. Due to the high mutational load from deposited Cas9, females developed as mosaic intersex individuals that are sterile. Mosaic males were selected for a downstream genetic cross (F2) to detect and assess generated mutations. **(B)** F2 male DsRed+YFP-offspring of the F1 cross were crossed to females containing a null allele of *dsx-F* marked by GFP^7^ (F2 cross). The GFP+ fraction of the offspring was selected using the COPAS fluorescent larval sorter, to balance mutations inherited from males. Females with intersex anatomy must contain non-restorative mutations (R2), and females showing normal anatomy must contain putative restorative (R1) mutations, or a wild-type unmodified allele. We analysed anatomically intersex mosquitoes *en masse* through pooled amplicon Illumina sequencing and anatomical females individually by Sanger sequencing. **(C)** The Sankey diagram shows the relative portion of GFP+ females that showed a female vs. intersex anatomy, and the relative proportion of the most common mutations discovered in them.

To maximise the number of *de novo* EJ alleles we performed 33 crosses of 50 F2 males each. To identify those alleles that may be resistant to gene drive activity, F2 males were crossed to an excess of females carrying a GFP-marked *dsxF*-null allele. Since *dsxF* is haplosufficient and the GFP-marked allele is null, we could differentiate those individuals that received a non-functional R2 mutation, and therefore displayed intersex phenotype, from those that received either an unmodified WT allele or an R1 mutation that restored DSX^F^ function (**Figure 1B**), and exhibited normal female anatomy. Assuming all 1650 males gave progeny, and distinct mutations were generated at the minimum rate of 2.75 per clutch, we calculated 4,538 *de novo* mutations that could be generated in this assay.

Among approximately 20,000 F3 progeny carrying the GFP-marked *dsxF*-null allele, that were examined as adults, 10,083 developed with non-male anatomy (i.e. female or intersex). The large majority (89.8%) of these non-male F3 displayed the full intersex phenotype (**Supp. Figure 6**), and pooled amplicon sequencing revealed that most of their alleles carried target site indels consistent with loss-of-function (**Figure 1C**, **Supp. Figure 8A, Supp. Figure 9**), most of which had been previously described^7^. The remaining 10.2% of non-male F3 comprised females with apparently normal external morphology, of which most were able to take a blood meal. Anticipating that putative R1 alleles could be enriched in these females, 852 were sequenced individually, revealing just 3.6% (n=31) that carried an unambiguous target site modification (**Figure 1B-C, Supp. Table 1**). Just 4 distinct alleles were identified in these individuals, all present in the blood fed group, and all causing an amino acid substation in the DSX^F^ protein. These putative R1 alleles include: a C→T substitution (2.2%) at position −3 of the T1 target site (2R:48714642, herein called T1-3C→T), a G→T substitution (1.1%) at position −4 of the T1 target site (2R:48714643, herein called T1-4G→T) and two alleles, containing more than 1 nucleotide substitution in the target site (0.4%) (**Figure 1C, Supp. Table 1, Supp. Figure 8B, Supp. Figure 10**). The two most common SNPs generated an alanine-to-valine and alanine-to-serine amino acid substitutions (**Supp. Figure 8B**).

We note that, occasionally, WT and putative R1 alleles were revealed at low frequency in the pooled sequencing of intersex individuals. This may be explained by ongoing stochastic mosaicism caused by vestigial Cas9 activity (zpg::Cas9 present in the *dsx*-mutator strain) in individuals that inherited the unmodified (WT) allele (see Methods, **Supp. Figure 5A, Supp. Figure 8A, Supp. Figure 9**).

### Natural and Cas9-induced functional SNP variants confer varying levels of resistance to the Ag(QFS)1 gene drive

Having identified both standing and drive-induced variants that are apparently functional, we next wanted to determine whether these variants are resistant to drive activity and therefore represent bona fide R1 resistant alleles. Therefore, we engineered *de novo* the most frequent natural variant (T1-2G→A at position 2R:48714641 in the genome, **Supp. Figure 4**), and the two most common putative **R1** variants identified in our mutagenesis screen (T1-3C→T and T1-4G→T, **Supp. Figure 8**) to confirm that they retain fertility and test whether they also confer resistance to the Ag(QFS)1 gene drive. The relevant nucleotide changes were precisely inserted into the genome using CRISPR-mediated cassette exchange (CriMCE), which allows the efficient recovery of specific defined mutations generated by HDR, even in the absence of a selectable marker (**Figure 2A-B**)^30^.

**Figure 2.**
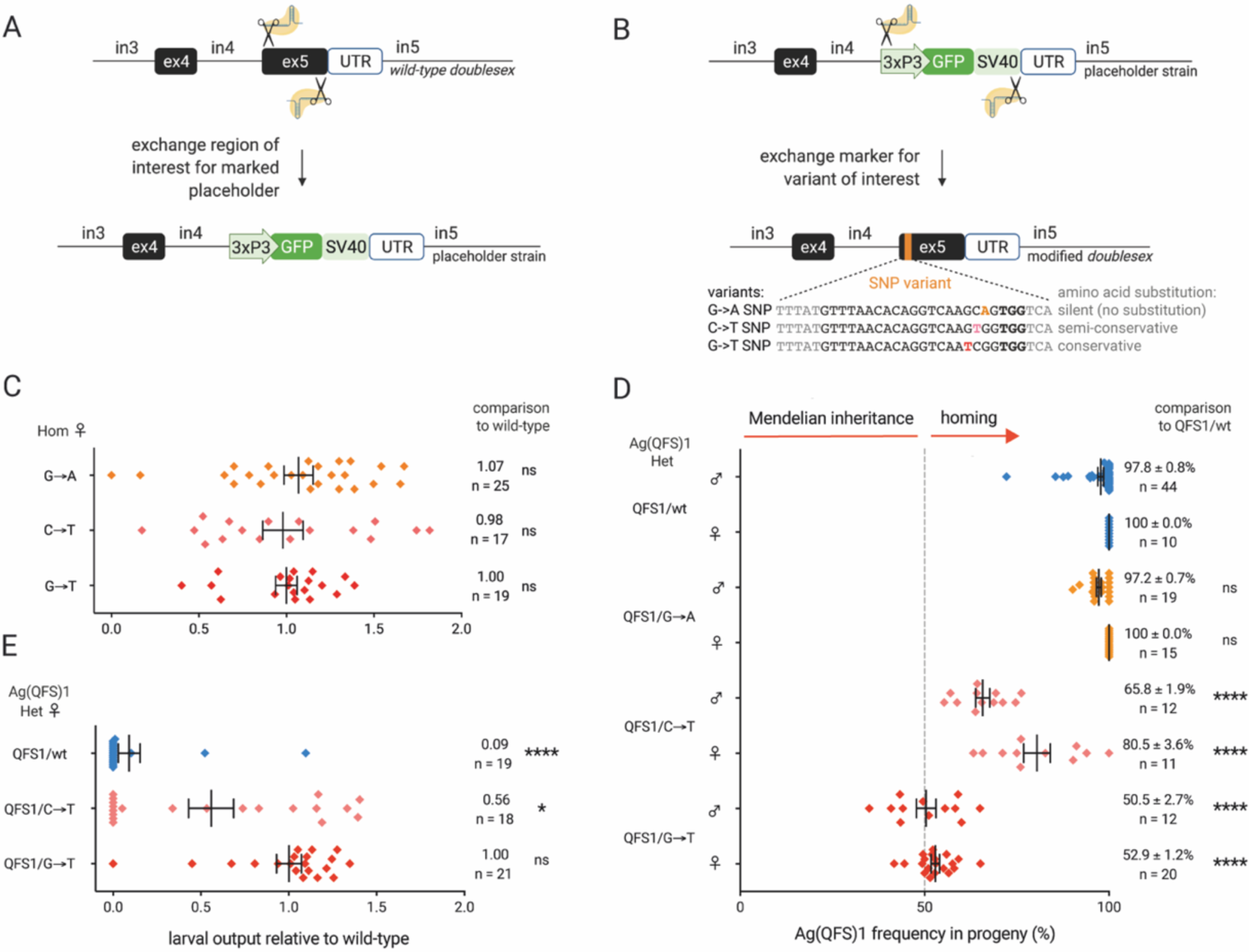
*De novo* engineering and testing of putative drive-resistant alleles revealed that each variant provided a different level of susceptibility to gene drive cleavage. **(A-B)** CRISPR-mediated cassette exchange (CriMCE) was used to engineer a naturally occurring (G→A, orange), and two Cas9-induced SNPs (C→T, pink and G→T, red) at the T1 gene drive target site on *dsx* exon 5. **(A)** First, the whole exon 5 was exchanged for a GFP-marked placeholder. **(B)** The placeholder was then exchanged for a version of exon 5 containing one of the SNPs of interest. **(C)** The relative larval output of homozygous (Hom) females for each SNP, compared to wild-type. Average larval output is shown to the right of each graph together with the sample size (n) and unpaired t-test statistical comparisons to each wild-type control (ns: non-significant with p=0.6459 for G→A, p=0.9946 for C→T, and p=0.4874 for G→T), after all datasets passed the D’Agostino & Pearson normality test. **(D)** Ag(QFS)1 frequency in the progeny of Ag(QFS)1/SNP trans-heterozygotes (Het), compared to Ag(QFS)1/wt. Mean Ag(QFS)1 transmission rates and the standard error around the mean (S.E.M.) are shown to the right of the graph, together with the sample size (n) and Kruskall-Wallis statistical comparisons to the Ag(QFS)1/wt control (ns: non-significant with p>0.9999, ∗∗∗∗: significant with p<0.0001). **(E)** The relative larval output of Ag(QFS)1/SNP and Ag(QFS)1/wt Het females, compared to a wild-type control. Mean relative larval output is shown to the right of each graph, together with the sample size (n) and non-parametric statistical comparisons to each wild-type control (Mann-Whitney:∗∗∗∗: significant, with p<0.0001 for Ag(QFS)1/wt, ns: non-significant, with p=0.0593 for Ag(QFS)1/C→T and p=0.7324 for Ag(QFS)1/GèT).

By performing blinded fertility assays we determined that females homozygous for each of the three engineered alleles are as fertile as wild-type (p=0.6459 for G→A, p=0.9946 for C→T and p=0.4874 for G→T) (**Figure 2C, Supp. Figure 11**). In homing assays, the T1-2G→A natural SNP was cleaved efficiently and did not reduce inheritance rates of the drive element in either sex (p>0.9999) (**Figure 2D**), and therefore does not represent a resistant allele, confirming our previous *in vitro* findings^7^. In contrast, the T1-4G→T SNP was found to be an **R1** allele that completely blocks the bias in gene drive inheritance rates in both sexes (p<0.0001) (**Figure 2D**). Interestingly, the T1-3C→T SNP did not block homing entirely, but significantly reduced drive transmission (65.8% inheritance in male carriers, p<0.0001; 80.5% inheritance in female carriers, p<0.0001) (**Figure 2D**). To our knowledge, this is the first description of a mutation that partially blocks gene drive activity, whilst maintaining gene function. We therefore termed this type of resistance **R3**.

Interestingly, the availability of the R1 allele allowed us to confirm a previous hypothesis that reduced fertility in female carriers heterozygous for the drive (Ag(QFS)1/wt) was due to ‘leaky’ Cas9 expression in the soma causing mutations in the WT allele^7,9,31^. Females heterozygous for the drive and the **R1** allele (Ag(QFS)1/R1) showed fertility equivalent to wild-type mosquitoes (p=0.7324), while those with the partially cleavable **R3** allele (Ag(QFS)1/R3) showed intermediate fertility (**Figure 2E**).

### The presence of partial resistance (R3) allows gene drive invasion but prevents population elimination

The litmus test of resistant alleles is their ability to be selected in a population and compromise the ability of a gene drive to invade. Starkly different population outcomes are possible depending on the type of resistance and the design of the gene drive itself^11,12,17,19^. Particularly interesting is the potential impact of **R3** alleles, which have not been described previously. Here, we introduced Ag(QFS)1 into caged populations pre-seeded with **R1** or **R3** resistance and tracked the frequency of resistant and drive alleles over time, together with the fraction of putatively fertile females as a measure of population suppression (**Figure 3A-B**).

**Figure 3.**
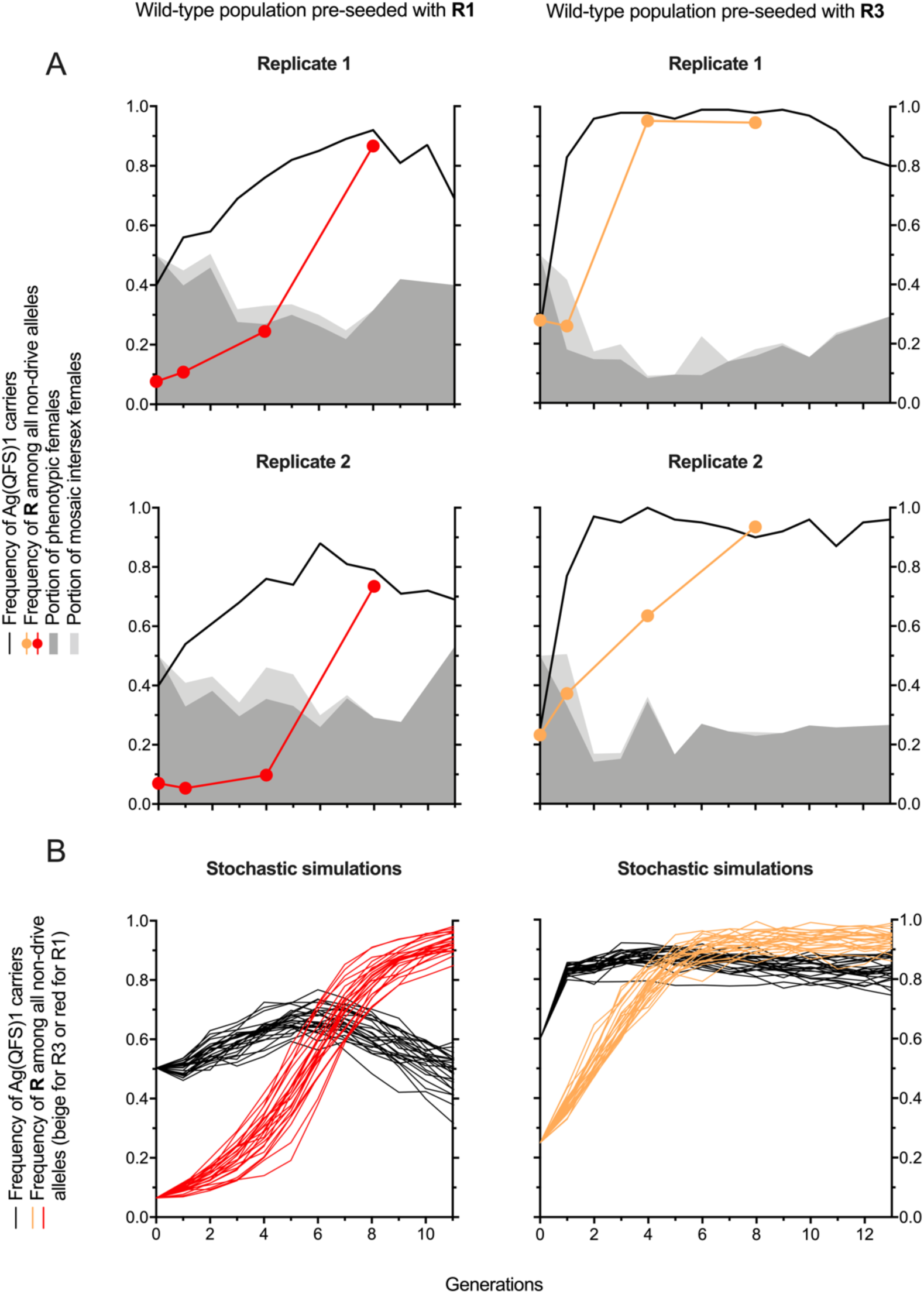
In the presence of R1 resistance Ag(QFS)1 spread is reversed, whereas in the presence of R3 resistance Ag(QFS)1 spreads, maintaining a high frequency, but ultimately failing to eliminate caged populations. **(A)** Ag(QFS)1 heterozygous males were released at a frequency of 40% (R1 cages, left) and 25% (R3 cages, right) in duplicate populations pre-seeded with 15% R1 (left) and 25% R3 (right) resistant alleles respectively. The frequency of Ag(QFS)1 carriers (solid black line) and resistant alleles amongst all non-drive alleles (red for R1 and beige for R3, circles denote sampled generations) was tracked over discrete generations using pooled amplicon sequencing. The proportion of females that developed normally as pupae (dark grey shaded region) or abnormally (mosaic intersex, light grey shaded region) is also shown. Note that fully intersex females and males are indistinguishable at the pupal stage from one another. **(B)** Stochastic simulations of caged populations pre-seeded with R1 (left) or R3 (right) resistance, using a fixed population size of 600 mosquitoes. Solid black lines refer to the frequency of Ag(QFS)1 carriers, and colour lines to the frequency of resistant alleles among all non-drive alleles (red for R1, beige for R3).

In cages seeded with both gene drive heterozygote males (at an overall frequency of 40%) and the **R1** allele (T1-4GèT) at a frequency of 9%, the **R1** allele increased rapidly within 8 generations, to 74-87% of all non-drive alleles in replicate cages (**Figure 3A**). In both replicates, as the **R1** allele began to reach high frequency the number of gene drive carriers started to decrease, in line with our predictions based on stochastic simulations (**Figure 3B**), and mirroring the dynamics observed for **R1** alleles at other gene drive target sites ^11,17^. As expected, in the presence of the **R1** allele there was little overall population suppression caused by the drive.

In cages seeded with gene drive heterozygote males (at an overall frequency of 25%) and the **R3** allele (T1-3CèT) at frequency of 25%, the observed invasion dynamics were markedly different to the case with the **R1** allele, and unique for a population suppression gene drive: near-full invasion of the drive did not cause complete population elimination, but rather a sustained ∼40% drop in reproductive output resulting from a reduction in the fraction of phenotypically normal females (**Figure 3A**). This is explained by the rapid selection of the **R3** allele that replaces the wild-type allele, resulting in a population segregating for it and an effectively weaker drive, meaning that though the drive spreads to near-fixation levels, it cannot eliminate the population – an outcome that is also predicted in stochastic modelling (**Figure 3B**).

To understand the potential impact of R1, R2 and R3 alleles, we simulated the dynamics of drive invasion and population suppression under different scenarios (**Figure 4**), assuming rates of allele creation determined in the mutagenesis screen (**Supp. Table 2**).

**Figure 4.**
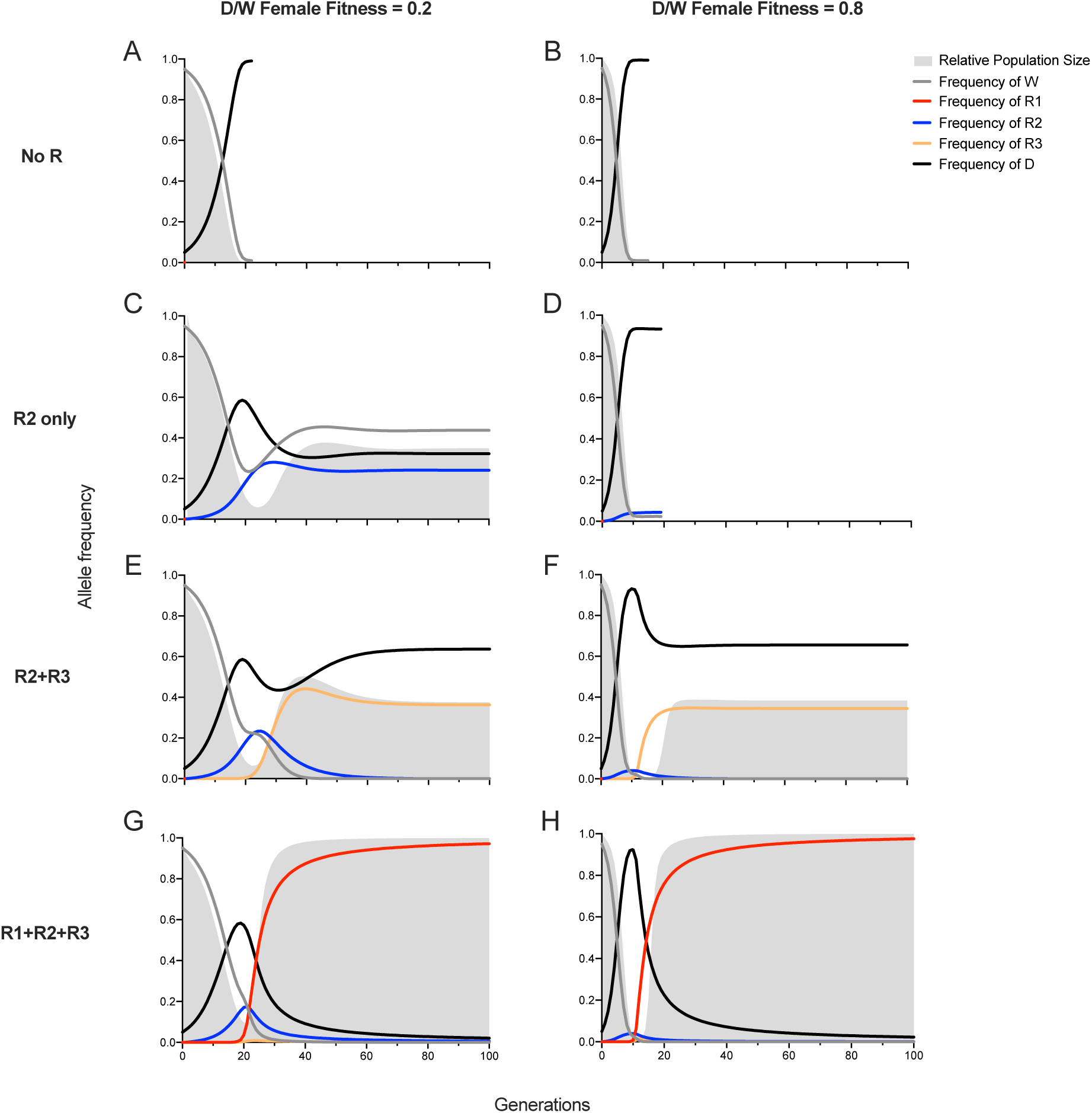
Effectively deterministic simulations of gene drive spread in the presence of no, R1, R2 and R3 resistance. W = wild-type, R1 = functional full resistance, R2 = non-functional resistance, R3 = functional partial resistance, D = gene drive, D/W = gene drive heterozygotes.

First, assuming no resistance, modelling simulations predict a rapid population elimination when the fitness of drive heterozygous females is high (fitness = 0.8), and similar but delayed dynamics when fitness is low (fitness = 0.2) (**Figure 4A-B**).

Second, assuming only non-functional **R2** resistant mutations arise, we observe rapid population elimination when the fitness of drive heterozygous females is high, but a stable suppression of ∼65% without elimination when fitness is low (**Figure 4C-D**). These results agree with previous findings^19^, supporting the hypothesis that population-level outcomes depend on the fitness of gene drive carriers.

Third, assuming partially resistant **R3** alleles arise in addition to **R2**, we predict a stable suppression without elimination (**Figure 4E-F**). In contrast to **R2**, the dynamics of spread and eventual stabilisation for **R3** are almost identical for both low and high fitness of drive heterozygous females, wherein **R3** alleles eventually displace both **R2** and wild-type. The extent of population suppression will depend upon the fitness and homing-blocking activity of the specific **R3** allele, which can be expected to vary greatly for different gene drive designs and resistant alleles. Assuming the characteristics of **R3** uncovered in this study, we predict a stable suppression of ∼60% (**Figure 4E-F**).

Finally, assuming all three types of resistant alleles arise (**R1**, **R2** and **R3**), we predict that the **R1** allele will become fixed in the population, displacing all other alleles including the gene drive, and allowing the population to recover in size (**Figure 4G-H**). These results are consistent with previous experimental and theoretical investigation of **R1** and **R2** resistance^11,17,19^.

### Multiplexed gene drives could mitigate resistance in natural populations

It has long been recognised that robust gene drive design should incorporate redundancy by targeting multiple sites simultaneously ^10,14,21^. To understand the potential effectiveness of this approach, we sought to estimate the rate of resistance at one target and extrapolate for a multiplexed gene drive assuming equal rates at all three sites.

Through our mutagenesis screen, we were able to estimate the rate (β or γ) at which either fully resistant (**R1**, β = 0.0016) or partially resistant (**R3**, γ = 0.0025) mutations might arise by EJ repair. In turn, using our stochastic modelling framework, we incorporated these values to calculate the mean probability of resistance arising in populations of different sizes (**Figure 5**). For comparison, we also calculated the probability of resistance arising at target sites with minimal sequence conservation, where a third of indels preserve the reading frame (**Figure 5**).

**Figure 5.**
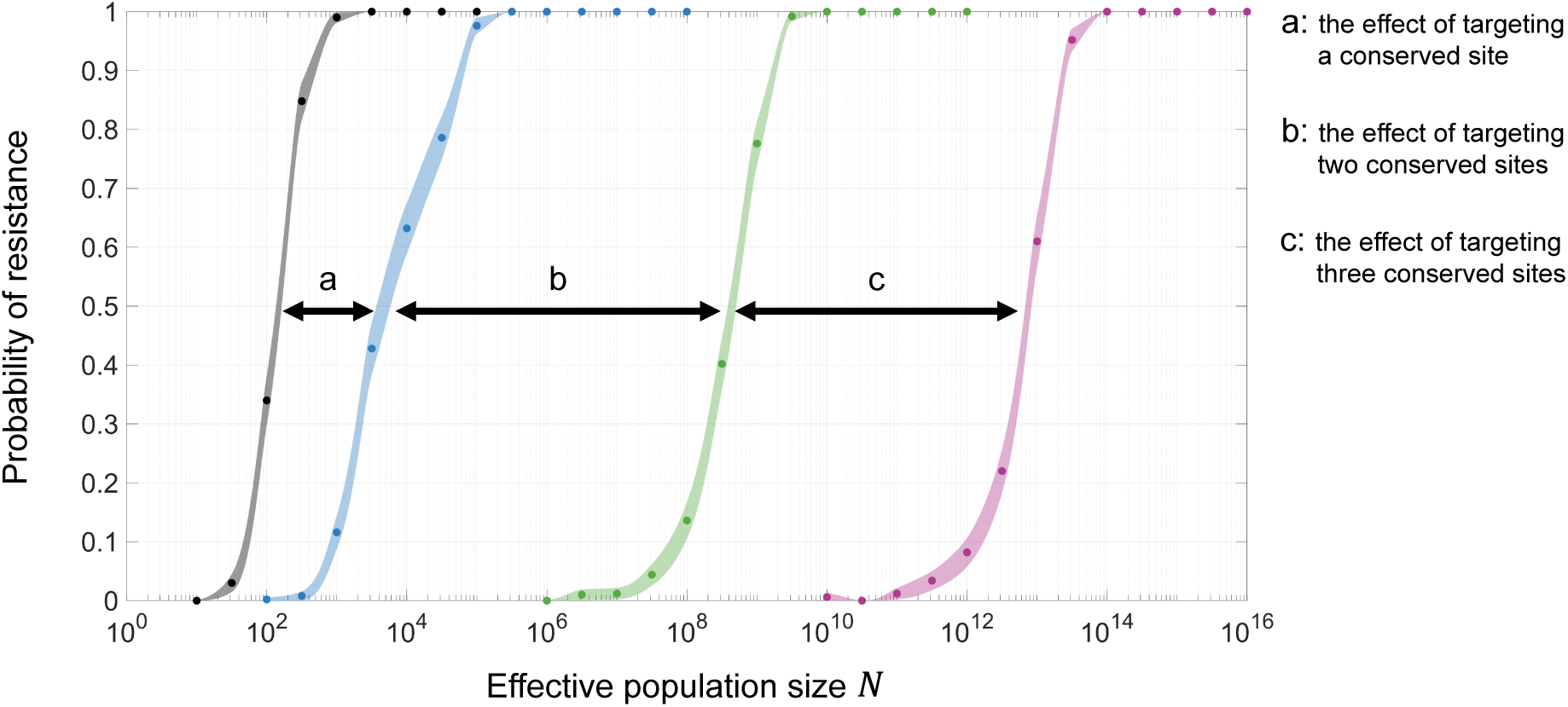
Simulations calculating the probability of resistance evolving against a gene drive targeting a single or multiple sites. Each data point is the average of 500 replicate stochastic simulations at the population size shown with 95% confidence intervals represented by the shaded regions, which have been smoothed for visual clarity. Simulations in grey assume a single gene drive target site with a rate of resistant allele creation = 1/3. Simulations in blue assume a single conserved gene drive target site, equivalent to the target site of Ag(QFS)1, with a rate of resistant allele creation as indicated by our resistance screen, i.e. β + γ = 0.0016 (for R1) + 0.0025 (for R3) = 0.0041. Simulations in green and violet assume the same rate of resistant allele creation for each target site, but for two or three target sites in total, respectively. For all simulations we assumed the same gene drive fitness as calculated for Ag(QFS)1. Resistance is defined by the population recovering to 95% of its original (wild-type) size.

Broadly, our results show that the probability of resistance increases monotonically as the population size increases, where the critical population size is roughly given by *N* × *β* ∼ 1, where *β* is the rate of generation of **R1** resistant alleles. We can define a maximum critical population size, *N_max_*, below which we can guarantee the prevention of resistance to at least 95% confidence. Firstly, we see that increasing the conservation of a site from *β* = 1/3 to *β* = 0.0016, increases *N_max_* from 38 individuals to 599 individuals (**Figure 5**). This is similar to the population size of existing population invasion experiments testing Ag(QFS)1 spread in small and large cages^7,9^. Indeed, simulations of laboratory experiments indicated that *de novo* generation of **R1** and **R3** is rare, with **R1** being expected to arise in 0.5% and **R3** in 1.9% of replicate cage experiments. Further, we predict that using 2 gRNAs would give protection up to *N_max_* ≈ 3.6 × 10^$^ individuals, whilst 3 gRNAs would give protection up to *N_max_* ≈ 5.2 × 10^%%^ individuals.

Estimates of natural mosquito population sizes vary between ≈ 10^&^ based on nucleotide diversity^27^, which is likely downward biased due to historical bottlenecks; to ≈ 10^$^ based on demographic history inference methods^32,33^; and up to ≈ 2 × 10^’^ based on the analysis of a recent soft sweep of insecticide resistance alleles, having updated our estimates given previously^34^, based on more recent estimates of the nucleotide mutation rate in *An. coluzzii* (*μ* ≈ 10())^35^. On this basis we would expect that 3 gRNAs targeting the female-specific exon of the *doublesex* gene (**Supp. Figure 3A**), would provide sufficient protection against resistance evolving.

### Multiplexing at the *dsx* locus improved drive transmission dynamics and mitigated resistance by actively removing resistant alleles

Based on our estimates of the perdurance of drives targeting multiple sites, we designed two new multiplexed gene drives to mitigate against the type of resistant alleles generated at the Ag(QFS)1 target site (T1). We developed two strains, targeting two (T1 and T3) or three sites on *dsx* exon 5 (T1, T2 and T3) (**Supp. Figure 3**), and named them Ag(QFS)2 and Ag(QFS)3 respectively (**Supp. Figure 12**). Both Ag(QFS)2 and Ag(QFS)3 gene drives showed high transmission rates that were comparable or higher than those of Ag(QFS)1 when targeting a wild-type locus (**Figure 6A-C, Supp. Figure 13**). This may be explained by overall higher cutting and repair at *dsx* in the presence of multiple guide RNAs^21^.

**Figure 6.**
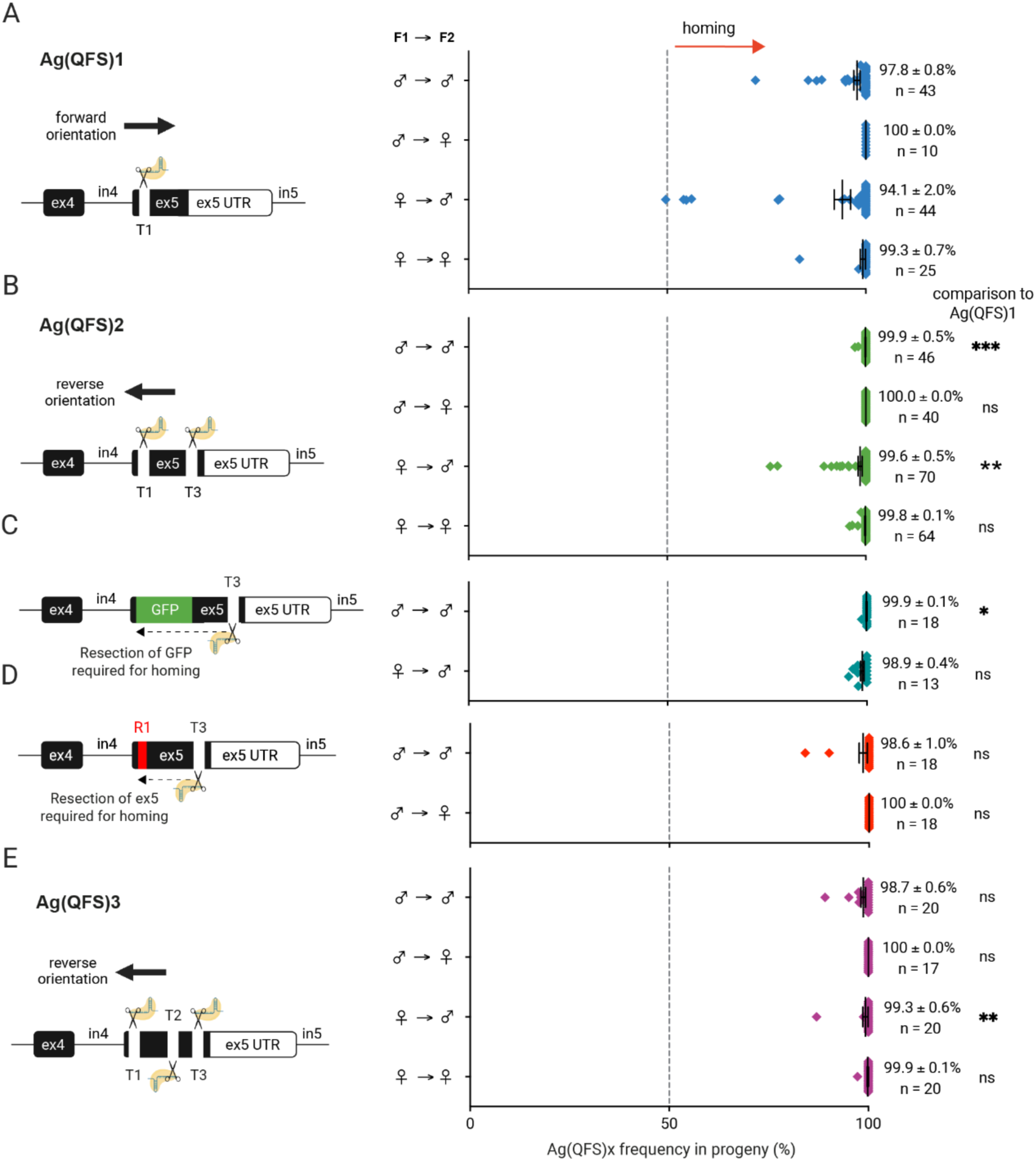
Transmission rates of multiplexed gene drives Ag(QFS)2 and Ag(QFS)3 in comparison to Ag(QFS)1. Schematics to the left show the gene drive tested, its orientation, and the number and location of available cut sites in each experimental set-up corresponding to results graphs to the right. Graphs to the right show the gene drive frequency in the progeny of gene drive heterozygotes: **(A)** Ag(QFS)1/wt (blue); **(B)** Ag(QFS)2/wt (green); **(C)** Ag(QFS)2/*dsxF-* (teal); **(D)** Ag(QFS)2/R1 (red); **(E)** Ag(QFS)3/wt (violet); each crossed to wild-type. The sex of the parents and grandparents of the scored progeny is shown to the left of each graph. For example, M→F indicates the frequency of the gene drive in the progeny of heterozygous gene drive females crossed to wild-type, that in turn inherited the gene drive from heterozygous gene drive males crossed to wild-type. Mean gene drive transmission rates and the standard error around the mean (S.E.M.) are shown to the right of each graph, together with the sample size (n) and Kruskall-Wallis statistical comparisons to the Ag(QFS)1/wt control (blue, panel A): **(B)** for Ag(QFS)2/wt, ∗∗∗: significant with p-value = 0.0004, ∗∗: significant with p-value = 0.0067, ns = non-significant with p-value > 0.9999; **(C)** for Ag(QFS)2/*dsxF-*, ∗: significant with p-value = 0.0206, ns = non-significant with p-value > 0.9999; **(D)** for Ag(QFS)2/R1, ns = non-significant with p-value > 0.1803 and p-value > 0.9999; **(E)** for Ag(QFS)3/wt, ns = non-significant with p-value > 0.9999, ∗∗: significant with p-value = 0.0057.

A fundamental tenet in employing a multiplexed drive is that it should still be able to home at a target locus containing at least one susceptible site; importantly, our multiplexed drive is designed such that, in the act of homing, it actively removes the resistant allele by forcing the resection and subsequent repair event beyond the resistant site (**Supp. Figure 2**).

Therefore, we tested the Ag(QFS)2 strain for its propensity to bias its inheritance when faced with the **R1** allele (T1-4GèT) at the T1 gene drive target site (**Figure 2D**). Ag(QFS)2 showed high transmission rates (98.6-100%) utilising exclusively the T3 site available for cleavage (**Figure 6D**). In this case a minimum of 78 nucleotides must be resected, past the **R1** allele, to generate ends with immediate homology to the drive allele to permit homing. In a separate experiment, forcing a resection of over 1kb, necessary to remove a dominant marker adjacent to a susceptible site, caused no detectable reduction in homing (**Figure 6C**). Together, these results suggest that there is ample width of sequence (at least 1kb) in which multiple sites could be chosen for a multiplexed gene drive, without affecting drive efficacy.

### Multiplexed gene drive carriers show marked improvements in fertility

Surprisingly, in characterising the multiplexed drive strains, in addition to the increased inheritance rates described above, we noticed that they also showed higher general fertility and reduced somatic mosaicism in heterozygous female carriers, compared to the original Ag(QFS)1 strain (**Figure 7A, Supp. Figure 14**). We still observed that mosaicism was higher in females inheriting the drive element paternally, rather than maternally, as we did for Ag(QFS)1^7,8^, although the effect was less pronounced (**Figure 7A-B, Supp. Figure 14**). We also observed increased post-bloodmeal mortality in Ag(QFS)1 heterozygous females, a phenotype that was also present, though, again, less pronounced, in Ag(QFS)2 heterozygotes (**Figure 7B**). Since blood-feeding is a female-specific trait in mosquitoes, it is perhaps not surprising that components relating to midgut metabolism might be controlled by DSX^F^.

**Figure 7.**
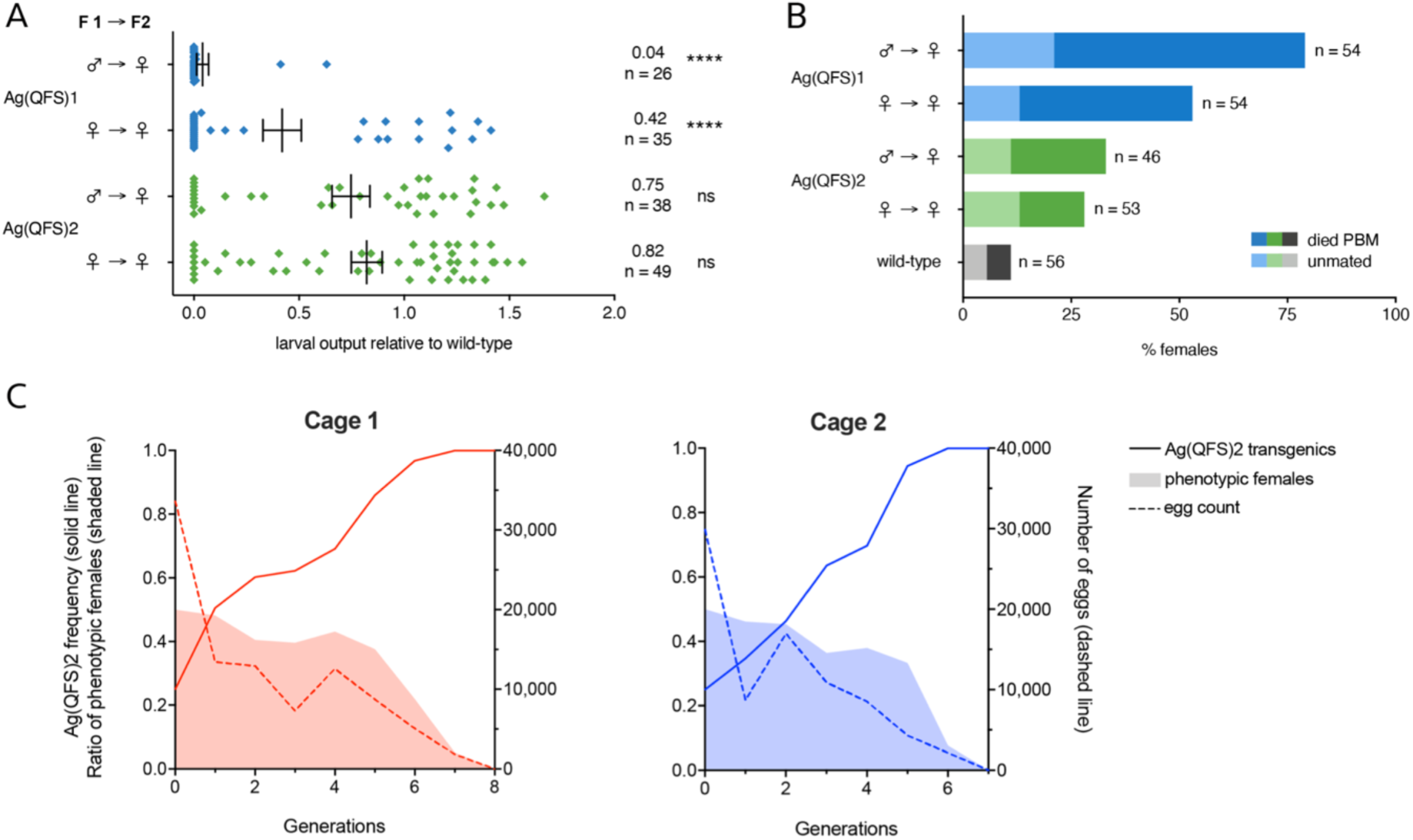
Improved fitness of Ag(QFS)2 multiplexed gene drive carriers contributes to efficient population invasion of small laboratory cages. **(A)** Female heterozygous Ag(QFS)1 or Ag(QFS)2 gene drive carriers that inherited the drive paternally (MèF) or maternally (FèF) were crossed to wild-type, and allowed to lay eggs individually. The same was done for a wild-type (WT) control. Egg and larval output of each female was recorded (**Supp. Figure 14**). Here we show larval output relative to the WT control. **(B)** The number of heterozygous females that died 3-4 days after receiving a bloodmeal was also recorded, as well as the mated status of live females. Wild-type females are included for comparison. Here, we show the percentage of unmated females (pale colours) and the percentage of females that died PBM (solid colours). **(C)** Ag(QFS)2 heterozygous males that inherited the gene drive paternally (MèM) were released in a wild-type population at a 25% (12.5% allelic frequency) in two replicate cages to make up a total population size of 600 for each cages that was maintained approximately constant over time by using 650 eggs of the previous to seed the following generation, until caged females were completely sterile and produced no eggs, at which point they were declared eliminated (at generation 8 for cage 1 and generation 7 for cage 2). The presence of Ag(QFS)2 (solid lines) was tracked over time through presence of the DsRed fluorescence marker. The proportion of phenotypic females was tracked over time as an indication of the reproductive capacity of the population (shaded regions), as well as the egg output of each cage (dashed line) per generation.

Of note, in the testing of the multiplexed Ag(QFS)3 lines, is that the level of somatic mosaicism differed markedly according to the orientation of the gene drive element integrated within the *dsx* target gene. When the transcriptional units controlling Cas9 and gRNAs were in the same direction as the target gene (‘forward’), levels of somatic mosaicism were about an order of magnitude higher in heterozygous carrier females than when the orientation of the drive element was in the opposite (‘reverse’) direction (74.9±13.9 % compared to 7.6±4.0%, respectively with p-value = 0.0104) (**Supp. Figure 15**). This effect was reproducible across Ag(QFS)3 lines generated independently from 5 different founder individuals. It therefore likely represents a genuine effect caused by the juxtaposition of certain elements within the drive element and the *dsx* target gene, rather than confounding effects such as deleterious alleles linked to the original integration event.

In light of this observation the improved performance of the multiplexed Ag(QFS)2 strain over the single guide Ag(QFS)1 may in part be attributable to the orientation of the drive element, the former being integrated in the reverse, and the latter in the forward orientation (**Supp. Figure 15**). Whether the effect of orientation is a direct consequence of the direction of transcription of the Cas9 or gRNAs relative to *dsx*, or of another feature related to the distinct gene drive architecture resulting from the two integration modes, is still unclear.

### Multiplexed gene drives targeting *dsx* can rapidly invade and supress laboratory populations

Taken together, multiplexed gene drive strains would be expected to show more efficient population invasion dynamics, even in the absence of any resistance. We therefore tested Ag(QFS)2 population invasion in laboratory cages, in a setup that mirrored the way we had originally tested the Ag(QFS)1 strain, using a release ratio of 1:1 heterozygous Ag(QFS)2 males to WT males (overall gene drive allele frequency of 12.5%)^7^. Ag(QFS)2 rapidly increased in frequency in two replicate populations, reaching complete fixation (100% gene drive carriers in the population) by generations 7 for cage A, and 6 for cage B, leading to a complete suppression in reproductive output by generations 8, and 7, respectively, due to the absence of fertile females (**Figure 7C**).

## DISCUSSION

Resistance poses an important challenge to the development and potential use of gene drive. New methods to mitigate resistance have allowed the successful modification of caged laboratory populations using gene drive, however bridging expectations between lab and field must anticipate a massive increase in population size. Here, we present a novel approach to assess the likelihood of resistance arising in large populations as an essential predictor of gene drive robustness and efficacy. Our strategy considers variants present in natural populations, and those generated through the mutagenic activity of Cas9. The improved throughput of our experimental set-up allowed us to show that one of the most functionally constrained gene drive target sites in the mosquito genome^22,27^ can evolve functional alleles, resistant to gene drive activity, where none had been detected before, in cage invasion experiments^7,9^.

We show that the only SNP found at appreciable frequency in natural populations is fully susceptible to Ag(QFS)1 gene drive activity, and therefore does not constitute a resistant allele. However, given the limited and non-uniform sampling of sequenced African malaria mosquito populations^26^, we cannot exclude yet-to-be detected resistant alleles residing at significant frequencies in the species.

We then applied a novel genetic screen to enrich for Cas9-induced mutations that encode a functional *dsx* gene product, recovering ∼4,000 mutations that may have arisen independently. This marks a dramatic increase in screening power over single generation gene drive crosses, or caged population experiments, where typically <1% of the individuals are expected to carry a target site mutation^7,9,17,36–38^. As an example, given the high homing rates for Ag(QFS)1 (93%, **Supp. Data File 1**), and EJ mutations being present in approximately half the non-drive progeny of heterozygote gene drive carriers^7^, over ∼500,000 gene drive offspring would need to be screened in order to have a power similar to our assay.

Through our genetic screen we discovered, and later validated, the first **R1** allele observed at the Ag(QFS)1 target site that restored full fertility to females and was fully resistant to gene drive activity. We show that when seeded into a caged population, this allele can prevent suppression by Ag(QFS)1, in agreement with previous research on **R1** resistance^11,17^. Additionally, we find an incomplete resistant allele, termed **R3**, that maintains *dsx* functionality but is only partially cleavable by Ag(QFS)1. Seeding lab populations with this allele did not prevent or reverse gene drive spread, but did prevent complete suppression. Indeed, our modelling predicts, that in the presence of **R3**, suppression drives will gradually level off to establish a stable equilibrium between gene drive and R3 alleles. This outcome is predicted to sustain a moderate level of long-term population suppression.

Importantly, our experimental approach allowed us to calculate the rate of creation of functionally resistant mutations (whether **R1** or **R3**) at 4.1*10^-3^ among all EJ alleles. Considering the rate of homing for Ag(QFS)1, the overall frequency among the offspring of gene drive carriers will be substantially lower at approximately 7.2*10^-5^ (**Supp. Data File 1**). At these rates, the likelihood of resistance to a gene drive like Ag(QFS)1, with only a single target site, exceeds 5% for populations greater than 600 individuals, and thus, it is a near certain outcome in natural populations.

Multiplexed gene drives are expected to reduce the likelihood of resistance by several orders of magnitude, if targeted to several non-overlapping sites showing strong sequence constraints ^36,38,39^. To this end, we designed and tested several multiplexed derivatives of Ag(QFS)1 and observed rates of homing at least equivalent to the single-target gene drive. In contrast to previous observations ^36,39^, we did not observe a reduction in homing when one of the target sites was blocked by the **R1** variant, or by an **R2** insertion (GFP) for which 1.4 kb of DNA resection would be needed to initiate homing. Essentially, the multiplexed design actively removes resistant variants if at least one site remains cleavable. Thus, multiplexing prevents population-level accumulation of single-site variants that can retard gene drive spread (**Supp. Figure 2**), and in doing so, also prevents the creation a fully resistant allele through the gradual accumulation of single-site mutations.

Consistent with previous observations at *dsx* and other gene drive targets^7,17^, we observed a fitness cost in Ag(QFS)1-heterozygous females that could be rescued when the gene drive was balanced against the **R1**. Unexpectedly, several multiplexed strains showed improved fitness linked to orientation of the drive elements relative to *dsx* – perhaps due to differences in somatic Cas9 expression.

Targeting several highly constrained sequences is expected to bring the rate of resistance below the threshold of detection in lab-contained releases. Our experimental approach allows such alleles to be created, and quantified, by first selecting for resistance at each site individually, and then combining variants onto a single synthetic, multi-resistant allele for testing. Assuming equivalent rates of resistance at each target site, our modelling determined that gene drives carrying 2 or 3 gRNAs could suppress effective population sizes of up to 3.6 × 10^7^ and 5.2 × 10^11^, respectively. Estimates of the effective population size of *An. gambiae* in Africa varies widely, from ≈ 10^7^ based on nucleotide diversity and demographic history inference methods^27^, to ≈ 2 × 10^8^ based on the analysis of a recent soft sweeps of insecticide resistance alleles^34^. This would indicate that 3 gRNAs would be sufficient to ensure protection against the evolution of resistance. However, there are many remaining uncertainties within these estimates as they assume a panmictic population^34^, ignore seasonality, and fail to consider complex spatial dynamics. Indeed, spatial models predict that persistent populations can arise under certain parameter conditions, where demographic stochasticity is enhanced^40–42^. Further modelling is needed to better predict heterogenous suppression and resistance over time.

## CONCLUSIONS

All suppressive technologies, from insecticides to antibiotics, are expected to exert evolutionary pressure for the selection of resistance against them. Since the inception of homing-based gene drive, theoretical exploration has considered different scenarios of resistance emerging, yet experimental evidence has been lacking^10,14,16^. By experimentally accelerating the emergence and selection of resistance, to the best-performing population suppression gene drive to date^7,9^, we have been able to make tangible and specific predictions around the scale and rate of resistance emergence at a highly constrained target - the female-specific exon of *dsx*. Such predictions, which are essential in moving from lab to field, were not possible with previous approaches. Finally, we demonstrate that multiplexed gene drives can actively remove resistant alleles, and when targeted to highly constrained loci such as *dsx*, are predicted to counteract resistance across large populations of the malaria mosquito. Our study should inform the future design of multiplexed gene drives to overcome resistance.

## MATERIALS & METHODS

### Analysis of standing variation at putative gene drive target sites

The Ag1000 phase 3 data release was accessed from Google Collaboratory, following instructions on the MalariaGEN website (https://www.malariagen.net/data/ag1000g-phase3-snp). The dataset includes genome-wide SNP calls from whole-genome sequencing of 2,784 wild-caught mosquitoes collected from 19 countries^26^.

### Mosquito maintenance and microinjections

*Anopheles gambiae* G3 strain wild-type and transgenic mosquitoes were reared at 26±2°C and 65±10% relative humidity. Larvae were maintained in trays with 500 ml salt water, at a larval density of 200 per tray and fed on NishiKoi food pellets. Adults were fed on glucose and females were blood-fed on cow-blood, 5-7 days after being crossed to males, using Hemotek membrane feeders^9^. Microinjections were performed on freshly laid embryos as previously described^7,43^. All plasmids were microinjected at a concentration of 300 ng/μl.

### Generation of an autosomal editor strain

To generate an autosomal editor strain expressing *zpg::Cas9* and U6::dsx-T1 gRNA and targeting the same site on *doublesex* (T1) as the Ag(QFS)1 gene drive we microinjected G3 strain wild-type embryos with the previously described p17410 plasmid^7^, together with a vasa-regulated *piggyBac* transposase helper plasmid^44^. The resulting strain is marked by *3xP3::DsRed* and expresses a U6::gRNA complementary to *dsx* T1 along with a *zpg::Cas9* that was insufficient in producing significant levels of *dsx* mutagenesis.

### High-throughput mutagenesis screen at *dsx*

To induce *dsx* T1 mutagenesis, an F1 cross of >100 heterozygous dsRed+ males carrying an autosomal editor expressing *zpg::Cas9* and U6::dsx-T1 gRNA, to >100 heterozygous *vasa2::Cas9* YFP+ females^4^ was performed 5 times, and females of each cross were blood-fed 5, 10 and 15 days after being crossed to produce offspring. 4 days post-bloodmeal (PBM) 4,000-12,000 L1 offspring of the F1 cross were screened and their DsRed+YFP-subsection was selected using a complex object particle analyser and sorter (COPAS)^45^. 50 DsRed+YFP-males were crossed to 50 GFP+ null heterozygous females (*dsxF-*)^7^ in an F2 cross. The F2 cross was performed 33 times to balance Cas9-induced mutations across the known null allele in their progeny and examine whether they can restore *dsx* functions in females. Each F2 cross was blood-fed once, and 4 days PBM 2,000-6,000 L1 offspring were screened and their GFP+ subsection was selected using COPAS^45^. GFP+ mosquitoes were reared to adulthood and 5 days post-emergence they were offered a blood-meal.

Subsequently, they were knocked-out using CO_2_ to examine their anatomy and manually sorted into 4 different pools per parental cage: males, intersex, blood-fed (BF) females, non-BF females. Intersex mosquitoes were identified by the presence of semi-plumose antennae and unrotated claspers^7^ (**Supp. Figure 6**). A pool of 100 intersex mosquitoes was analysed from each parental cage by pooled amplicon sequencing. Mosquitoes that developed normally as females were individually analysed by Sanger sequencing. Note that samples were individually analysed several months after being collected due to interruption of the study by the Covid-19 pandemic.

### Pooled amplicon sequencing

Pools of adult mosquitoes (100 intersex mosquitoes per pool for each of the resistance assay cages and ∼250 mosquitoes in duplicate per pool for each of the cage trial generations) were subjected to gDNA extraction using the Promega Wizard® Genomic DNA Purification kit, PCR amplification under non-saturating conditions as previously^7,17^, using primers Illumina-AmpEZ-4050-F1 (<u>ACACTCTTTCCCTACACGACGCTCTTCCGATCT</u>ACTTATCGGCATCAGTTGCG) and Illumina-AmpEZ-4050-R1 (<u>GACTGGAGTTCAGACGTGTGCTCTTCCGATCTGT</u>GAATTCCGTCAGCCAGCA), Illumina pooled amplicon sequencing (AmpEZ service, Genewiz, Azenta Life Sciences) and analysis using CRISPResso2^46^. Datasets were subsequently handled on Python to rename mis-labelled alleles that were falsely grouped together by CRISPResso. A detection threshold for mutations was set at 0.5%.

### Sanger sequencing

Single mosquito samples were subjected to gDNA extraction using the DNeasy Blood & Tissue kit (Qiagen), PCR amplification using primers dsx-intron4-F1 (GTGAATTCCGTCAGCCAGCA) and dsx-exon5-R4 (AACTTATCGGCATCAGTTGCG), and Sanger sequencing by Eurofins Genomics using primer dsx-exon5-R2 (TGAATTCGTTTCACCAAACACAC). Sequence alignments were performed on Benchling.

### Generation and phenotype assessment of the *dsx* variant strains

The three marker-less variant strains containing different SNPs at the *dsx* T1 site (GèA, CèT or GèT) were generated and isolated using CriMCE as previously described^30^. Female fertility of SNP homozygotes and heterozygotes was determined by double-blinded assays, whereby mixed populations of wild-type and SNP carrier females were crossed to wild-type males and allowed to lay eggs individually 2-3 PBM. Eggs and larvae were counted no more than 1 day after they were laid or hatched respectively. The genotype of each mother was then determined by Sanger sequencing (wild-type, heterozygous or homozygous for each SNP). To determine the extent to which each SNP was cleavable by the gene drive, SNP carrier females were crossed to DsRed+ Ag(QFS)1 gene drive males and their trans-heterozygote Ag(QFS)1/SNP offspring, as well as Ag(QFS)1/wt individuals of both sexes, were crossed to wild-type. Females were allowed to lay eggs individually 2-3 days PBM. The rate at which Ag(QFS)1 was inherited amongst the offspring was determined through fluorescence microscopy by tracking the DsRed marker. The egg and larval output of Ag(QFS)1/CèT, Ag(QFS)1/GèT and Ag(QFS)1/wt females was also recorded and compared to a simultaneous wild-type control, respectively.

### Population invasion experiments

Population invasion experiments were set up by seeding wild-type populations with either **R1** or **R3**, and the Ag(QFS)1 gene drive. Briefly, **R1**-seeded populations were setup in duplicate cages, initiated with 240 Ag(QFS)1 heterozygous gene drive males (40% genotype frequency) and a mixed population of 306 wild-type and 54 **R1** homozygotes (9% allelic frequency). **R3**-seeded populations were setup in duplicate cages, initiated with 150 Ag(QFS)1 heterozygous gene drive males (25% genotype frequency) and a mixed population of 450 mosquitoes carrying **R3** at approximately 25% allele frequency (the progeny of **R3** heterozygous females crossed to wild type).

The populations were mixed as pupae in the starting generation (G0) and were allowed to emerge in the same cage. Adult mosquitoes were left to mate for 5 days post-emergence, before being blood-fed on cow blood using a Hemotek membrane feeding system for 30 minutes. Filter paper-lined cups of water were placed in the cages 2 days PBM. Eggs were collected and photographed 3 days PBM. 1^st^ instar larvae (L1) were screened 4-5 days PBM, by tracking the DsRed fluorescence associated with the gene drive through fluorescence microscopy or by using the COPAS larval sorter^45^. A minimum of 600 larvae were randomly selected and split into 3 trays of 500ml water (200 larvae per tray). All surviving pupae were collected and transferred to a new cage for the adults to emerge and establish the following generation. A representative portion of these (100-300 individuals) were screened as pupae to determine the sex ratio per generation. Note that males and intersex are indistinguishable at the pupal stage, however mosaic intersex females and phenotypically typical females can be distinguished. After blood-feeding and oviposition of each following generation the adults of the previous generation were collected and stored in the −20°C for further analysis by pooled amplicon sequencing to determine the frequency of **R1** and **R3** alleles in each cage.

### Modelling

We perform two types of simulations: (i) effectively deterministic simulations that include a partial resistance allele **R3** with Beverton-Holt population dynamics (**Figure 4**) and (ii) stochastic simulations of cages that are maintained at a fixed (**Figure 3B**) or varying population size (**Figure 5**). Both use the same underlying stochastic modelling framework, as previously described^16^, and are based on a Wright-Fisher model for population genetics with two separate sexes. For simulations of type (i), we use a very large initial population size (*N_e_* = 10^%+^) that mimics deterministic dynamics, which use a very accurate and computationally efficient approximation to multinomial sampling^16^; however, as a result of the population dynamics, when the total population size is small this results in stochastic effects just before population elimination, as such in the instances where the population size got drastically reduced and the populations eliminated, we have only plotted allele frequencies for generations in which *N_e_* > 10,000. For simulations of type (ii), we use a fixed small population size of *N_e_* = 600 (**Figure 3B**) or varying population sizes from *N_e_* = 10^0^ up to *N_e_* = 10^16^ to determine the critical population size that can be eliminated by a gene drive with a single, two or three target sites with 95% confidence (**Figure 5**). For simplicity, a standard fitness cost of 0.76 (i.e. the fitness cost of Ag(QFS)1 heterozygous females) was applied for any female gene drive carrier, since fitness only impacted upon the critical population size calculation for a single-target gene drive, but not for a gene drive containing two or three target sites. To account for the observed initial advantage of Ag(QFS)1 males in the experimental **R3** cages, due to the separate rearing of Ag(QFS)1 and wt/R3 individuals, we started the initial Ag(QFS)1 frequency at 0.6 in the simulations to approximately match the cage data (**Figure 3B**).

The key difference between the simulations in this paper and previous modelling of resistance to gene drive^16^, is the inclusion of the partially resistant allele **R3**. The process of gametogenesis modelled in the simulations is shown in **Supp. Figure 9**, with parameters as described in **Supp. Table 2** and **Supp. Figure 16**. Note that the only genotypes are inherited in a super-Mendelian fashion are W\D (Ag(QFS)1/wt) and R3\D (Ag(QFS)1/R3). We assume that **R1** and **R2** alleles completely block gene drive homing.

### Cloning of CRISPR constructs with multiplexed gRNAs

Primers containing *Bsa*I sites (capitals) and gRNA sequences (capitals): BsaI-T1-U6-F (gagGGTCTCatgctGTTTAACACAGGTCAAGCGGgttttagagctagaaatagcaagt) and BsaI-T3-U6-R (gagGGTCTCaaaacCTCTGACGGGTGGTATTGCagcagagagcaactccatttcat), were used to amplify a gRNAscaffold-U6terminator-U6promoter cassette from plasmid p131^30^. The resulting PCR product was inserted into the p174 master-vector previously described^7^, through GoldenGate cloning to create vector p174102 that was used as a helper plasmid to generate the Ag(Dsx) docking line, and as a donor vector to generate the Ag(QFS)2 gene drive. This vector (p174102) contains a *zpg::hCas9* nuclease cassette, *3xP3::DsRed::Sv40* marker, and two tandemly repeated *U6::gRNA* cassettes targeting sites T1 and T3 on *dsx* exon 5 (**Supplementary Figure 7**). To create the p174103 vector for generation of the Ag(QFS)3 strain, oligonucleotide sequences: dsx-T2-F (TGCTGTCTGAACATGCTTTGATGGCG) and dsx-T2-R (AAACCGCCATCAAACATGTTCAGAC) were annealed and cloned into the p139 vector containing a gRNAscaffold-U6terminator-U6promoter-BsaIcloningsite-gRNAscaffold-U6terminator-U6promoter cassette, through GoldenGate cloning. The resulting plasmid was amplified using primers BsaI-T1-U6-F and BsaI-T3-U6-R like before, to enable GoldenGate cloning of the PCR product into p174, to create p174103, containing a *zpg::hCas9* nuclease cassette, *3xP3::DsRed::Sv40* marker, and three tandemly repeated *U6::gRNA* cassettes targeting sites T1, T2 and T3 on *dsx* exon 5 (**Supplementary Figure 7**).

### Cloning of donor plasmids to generate the Ag(Dsx) gene drive docking line

The pDLv3 donor vector, used for generation of the Ag(Dsx) strain, was created by Gibson assembly. A 3xP3::GFP::SV40 marker cassette flanked by *attP* sites was amplified using primers 4050-KI-Gib34 (ATGTTTAACACAGGTCAAGACCGGTACCCCAATCGTTCA) and 4050-KI-Gib35 (GCGGAAAGTTTATCATCCACTCACGCGTTCAGGATTATATCT). Genomic regions 1.8 kb upstream and downstream of the dsx intron4-exon5 splice junction were amplified using primer pairs 4050-KI-Gib1 (GCTCGAATTAACCATTGTGGACCGGTCTTGTGT TTAGCAGGCAGGGGA) with 4050-KI-Gib33 (TGAACGATTGGGGTACCGGTCTTGACCTGTGTTAAACATAAATG), and 4050-KI-Gib36 (AGATATAATCCTGAACGCGTGAGTGGATGATAAACTTTCCGCAC) with 4050-KI-Gib4 (TCCACCTCACCCATGGGACCCACGCGTGGTGCGGGTCACCGAGATGTTC), to make up the left and right homology arms of the donor plasmid, respectively. The pK101 plasmid^7^ was digested with restriction enzymes *Mlu*I and *Bsh*TI to release its 5.4 kb backbone containing a 3xP3::DsRed::SV40 marker, which was combined with the three amplicons described, in a 4-fragment Gibson assembly reaction, so that the *dsx* homology arms get placed to each side of the *attP*-flanked eGFP cassette.

### Generation of the Ag(Dsx) docking strain

To generate the Ag(Dsx) docking line to accommodate recombinase-mediated cassette exchange (RMCE) of plasmids p174102 and p174103 to generate the Ag(QFS)2 and Ag(QFS)3 gene drive constructs, respectively, wild-type embryos of the *Anopheles coluzzii* G3 strain were microinjected with the p174102 CRISPR plasmid and the pDLv3 donor plasmid. In transformants, CRISPR elements expressed by p17402 caused excision of the region between the T1 and T3 cut sites on *dsx* exon 5, and its replacement with the *attP*-flanked GFP marker cassette from the pDLv3 plasmid, facilitated by HDR. All microinjection survivors (G0) were outcrossed to wildtype mosquitoes and positive transformants were identified through fluorescence microscopy as GFP+ and RFP-.

### Generation of the Ag(QFS)2 and Ag(QFS)3 multiplexed gene drive strains

To generate the Ag(QFS)2 and Ag(QFS)3 multiplexed gene drive strains, heterozygotes of the Ag(Dsx) strain were crossed to each other, and their progeny were microinjected with the p174102 or p174103 CRISPR plasmids, respectively, together with a vasa-integrase helper plasmid^44^, to facilitate RMCE of the GFP cassette of Ag(Dsx) for each of the gene drive constructs. All microinjection survivors (G0) were outcrossed to wild-type mosquitoes and positive transformants were identified through fluorescence microscopy as dsRed+ and GFP-. Construct orientation was determined via PCR using primer pairs zpg-term-F1 (GCTGTACTACATCTCGTGGACG) with dsx-exon5-R2, which would only produce an amplicon if the gene drive construct is integrated the same orientation as the *dsx* gene (fwd), and zpg-term-F1 with dsx-intron4-F1, which would only produce an amplicon if the gene drive construct is integrated in reverse (rev) with respect to *dsx*.

### Phenotype assessment of multiplexed gene drive strains

Gene drive homing of strains Ag(QFS)2 and Ag(QFS)3 was assessed by crossing heterozygous dsRed+ gene drive parents of both sexes that inherited the drive from either males or females to wild-type. Ag(QFS)2 male individuals that inherited the gene drive from either males or females, were also crossed to heterozygous females of a GFP+ *dsxF*-strain^7^, where a GFP cassette interrupts target site T1 but leaves T3 exposed to cleavage.

Trans-heterozygous GFP+ dsRed+ offspring of both sexes were crossed to wild-type to determine rates of gene drive inheritance in the offspring. After allowing females to lay eggs individually, 2-3 days post-bloodmeal, the rate at which each gene drive (DsRed+) was inherited amongst their progeny was determined through fluorescence microscopy. The fertility of Ag(QFS)1, Ag(QFS)2 and wild-type females was compared through fertility assays performed simultaneously to allow direct comparison. These looked at egg and larval output, mating ability and post-bloodmeal mortality. Females that inherited each drive from either males or females, as well as wild-type females, were crossed to wild-type males, and allowed to lay eggs individually 2-3 days post-bloodmeal. Their egg and larval output were counted no longer than 1 day post laying and hatching, respectively. The number of females that had died in any of the 3-4 days post-bloodmeal were recorded.

Females were also interrogated for their mating ability by examining their spermathecae for sperm presence under an EVOS high resolution light microscope. Note that previously, phenotypic assays of Ag(QFS)1 excluded from the analysis females that were unmated and females that did not blood-feed ^7^. In the present study all females were included in the analysis to estimate a more accurate measurement of female fertility, since blood-feeding and mating ability might be affected by the *dsx*F knockout. Finally, the number of females showing a mosaic intersex phenoype amongst progeny of males of different gene drive strains that were integrated in forward (fwd) or reverse (rev) orientation with respect to the dsx gene, were counted. This included Ag(QFS)1 (fwd orientation), Ag(QFS)2 (rev orientation) and five independently created Ag(QFS)3 strains of either orientation.

### Population invasion experiments of Ag(QFS)2 in small cages

Population invasion experiments were performed in duplicate with each G0 population consisting of 300 wild-type females, 150 wild-type males and 150 Ag(QFS)2 DsRed+ heterozygous males that inherited the gene drive paternally, as previously^7^. The initial caged release was performed using age-matched pupae. Mosquitoes were left to mate for 5 days post-emergence before being blood-fed on cow-blood. Two days later they were allowed to lay eggs. 650 eggs were randomly selected to seed the next generation, whilst the rest were photographed and counted using JMicroVision. Emerged larvae were screened for the presence of DsRed, indicative of the gene drive, using fluorescence microscopy. After pupation the percentage of phenotypic females amongst all pupae including males/intersex and mosaic intersex, was recorded to give an indication of the reproductive capacity of the caged populations, before allowing all pupae indiscriminately to seed the next generation. Note that sterile intersex mosquitoes are indistinguishable from males at the pupal stage.

### Data visualisation

Graphs were plotted on Graphpad Prism 9, and figures were designed on Biorender (full licence), Sankeymatic (https://sankeymatic.com/), MapChart (https://www.mapchart.net/) and Adobe Illustrator.

## Author Contributions

I.M.: Conceptualisation, Methodology, Experimental Work, Data Analysis, Data Interpretation, Writing – Original Draft, Writing – Review & Editing, Data Visualization. L.P.: Experimental Work. B.S.K.: Mathematical Modelling, Data Visualization, Writing – Review & Editing. L.M.: Experimental Work. M.G.: Experimental Work. A.B.: Writing – Review & Editing, Funding Acquisition. F.B.: Writing – Review & Editing, Project Administration. A.M.H.: Conceptualisation, Data Interpretation, Writing – Original Draft, Writing – Review & Editing, Supervision. T.N.: Conceptualisation, Data Interpretation, Writing – Original Draft, Writing – Review & Editing, Supervision, Project Administration. A.C.: Conceptualisation, Supervision, Funding Acquisition. All authors approved the manuscript.

## Acknowledgements

We thank Kyros Kyrou for data sharing and useful discussions; Nace Kranjc for bioinformatic assistance; Molly McGrath for her assistance in molecular cloning; John Connolly, Silke Fuchs and Alekos Simoni for providing useful comments on the manuscript draft; and the whole of Crisanti Lab for overall support, especially during the Covid-19 pandemic.

---

**Supplementary Figure 1.**
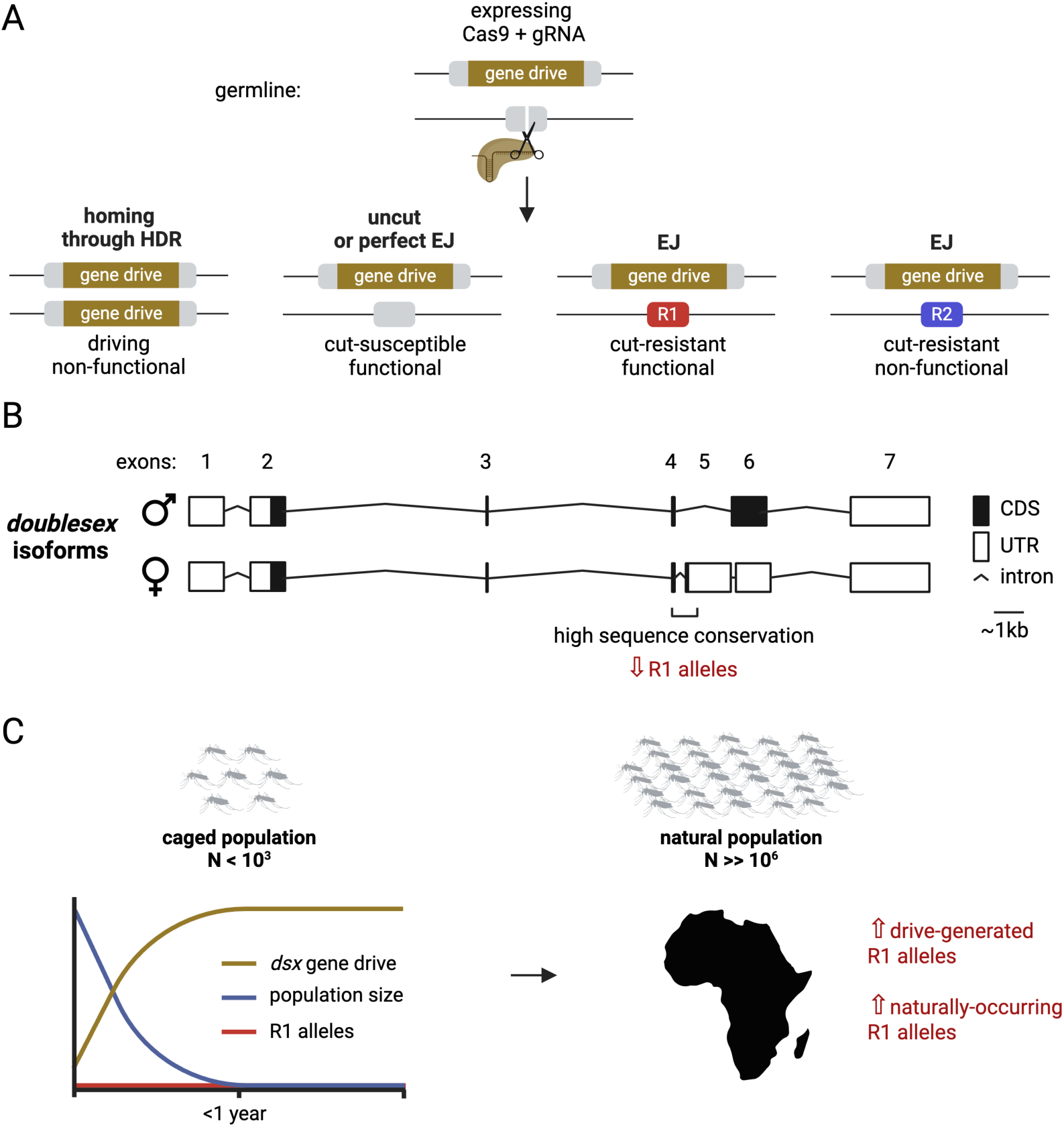
The generation of drive-resistant mutations. **(A)** There are four main repair outcomes after a gene drive-derived Cas9 and gRNA catalyse cleavage of an exposed chromosome, carrying their intact target site, in the germline. The most common outcome is homing of the gene drive (light brown) due to homology-directed repair (HDR). Alternatively, the cut chromosome gets repaired by end-joining (EJ). Perfect EJ repairs the wild-type allele (light grey), which is susceptible to further cleavage. EJ is often error-prone and can lead to the introduction of cut-resistant mutations at the gene drive target site that are either functional (R1, red) or non-functional (R2, blue). Functional resistance (R1) has a selective advantage and can reverse gene drive spread. **(B)** The *doublesex* gene is expressed into two sex-specific isoforms. Targeting the highly conserved, and presumably functionally constrained region on the intron-exon boundary of the female-specific exon (exon 5) of *dsx*, can limit R1 alleles. **(C)** Indeed, no R1 alleles were detected in caged laboratory populations, leading to complete population elimination in less than a year^7,9^. However, natural populations are larger, by several orders of magnitude, which increases the likelihood of drive-induced R1 allele formation, whilst R1 alleles might also be pre-existing in nature.

**Supplementary Figure 2.**
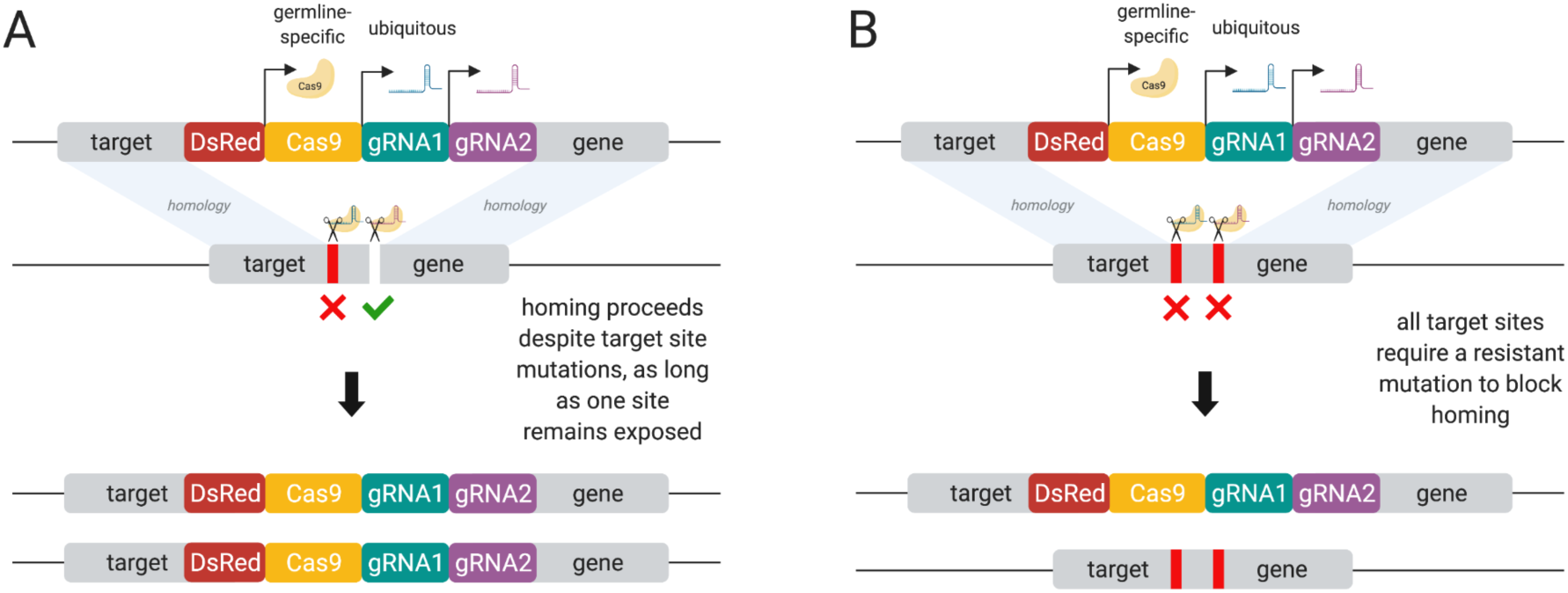
Multiplexed gene drives can mitigate resistance by targeting multiple sites simultaneously. **(A)** Resistant mutations (shown in red) are removed from the target locus provided that one of the targeted sites remains cleavable, to permit homing. **(B)** To block gene drive homing all target sites would need to carry a resistant mutation (shown in red).

**Supplementary Figure 3.**
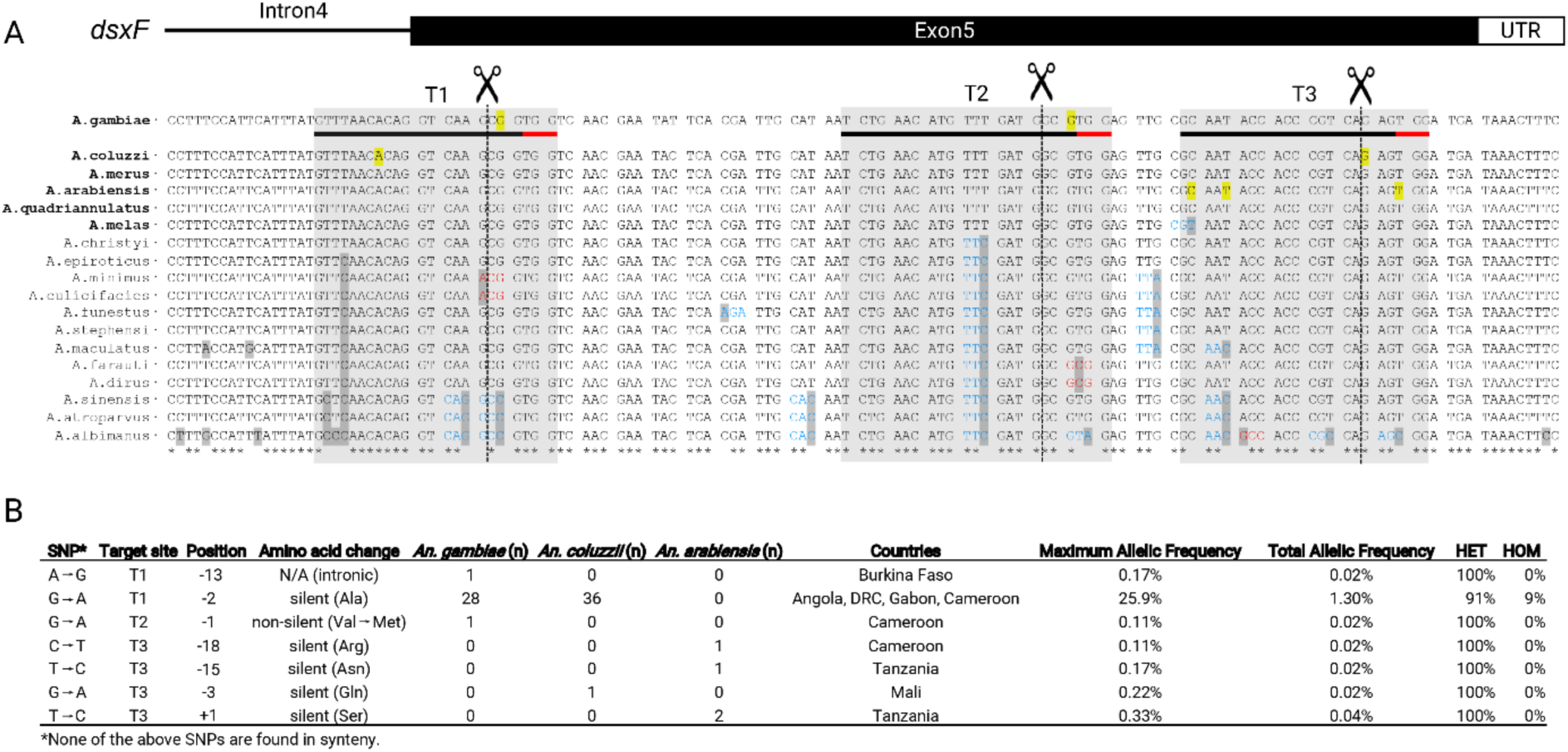
Natural variation on the coding sequence of the female-specific exon of *doublesex*. **(A)** Multiple *Anopheles* species alignment reveals a high amount of nucleotide conservation in both the coding region of exon 5 and flanking non-coding regions (intron 4, exon 5 UTR). In bold are species that belong to the *An. gambiae* species complex. Nucleotide changes are highlighted in grey and those leading to amino acid changes are in red, whilst silent changes are in blue. Three sites that could be targeted by gene drive (T1, T2 and T3) are shaded and underlined on the *An. gambiae* reference. Variable nucleotides within them, as identified by analysing the phase 3 Ag1000G project data are highlighted in yellow. Their proto-spacer adjacent NGG motifs (PAM) are underlined in red. The dashed line pinpoints the cleavage sites. **(B)** The table shows single nucleotide polymorphisms (SNPs) that were identified amongst 2,784 wild-caught mosquitoes in *An. gambiae, coluzzii* or *arabiensis*, and their relative positions compared to each cleavage site in T1, T2 or T3. Maximum allelic frequency refers to the frequency of each SNP in the country in which it was most frequent. The percentage of mosquitoes that possessed each SNP in heterozygosis (HET) vs homozygosis (HOM) is also shown.

**Supplementary Figure 4.**
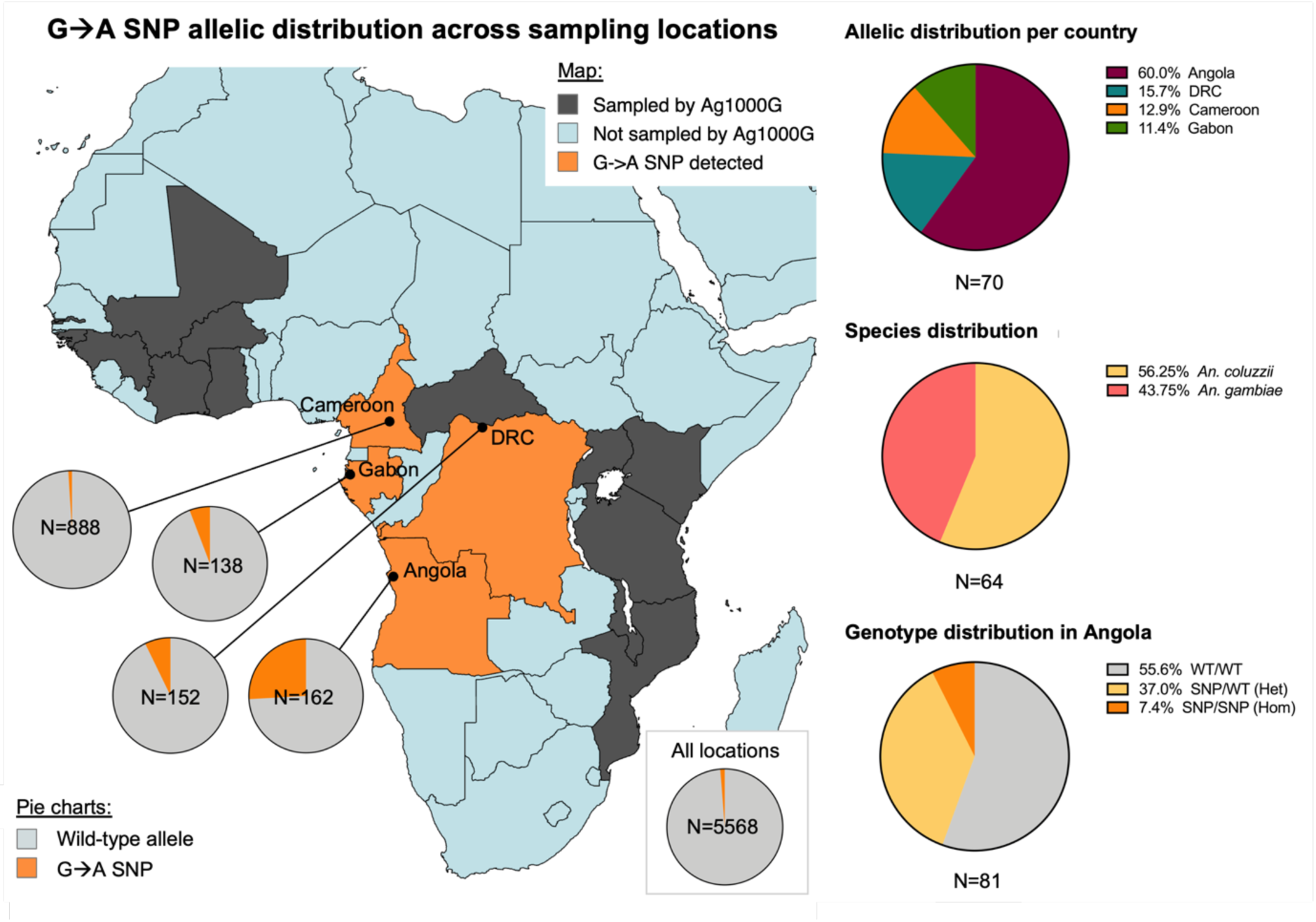
The allelic distribution of the natural G→A variant (2R:48714641), across Ag1000G sampling locations. The G→A natural variant was found at 1.3% allelic frequency across all sampling locations, including 19 African countries and territories (N=5568 alleles). Specifically it was found at 1.0% allelic frequency in Cameroon (N=888 alleles), 5.8% allelic frequency in Gabon (N=138 alleles), 7.2% allelic frequency in the DRC (N=152 alleles) and at 25.9% allelic frequency in Angola (N=162 alleles). The SNP’s allelic distribution per country is shown to the top right (N=70 alleles), its species distribution to the middle right (N=64 alleles), and its genotype distribution in Angola (N=81 genotypes), where it was most frequent to the bottom right. The map of Africa was designed using MapChart (https://www.mapchart.net/).

**Supplementary Figure 5.**
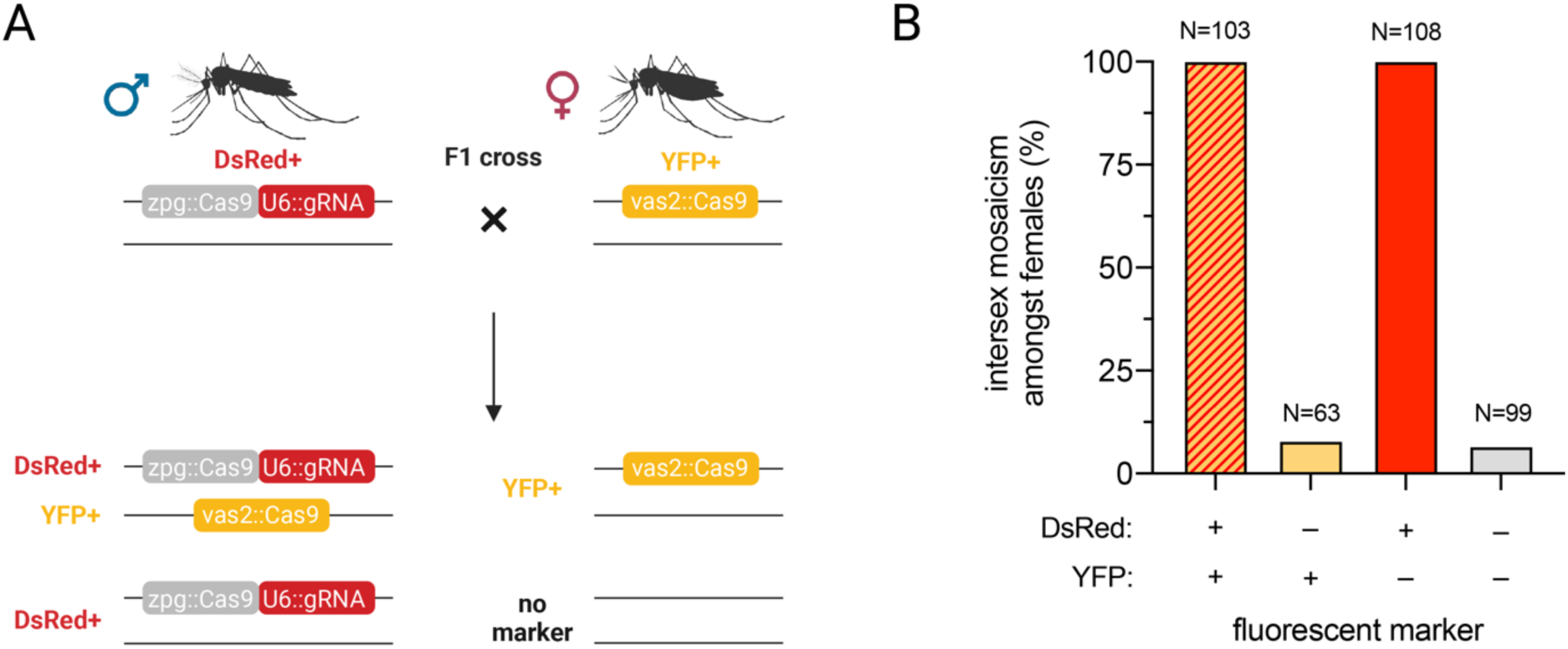
By exploiting maternal deposition of Cas9 we can generate high rates of end-joining mutations at the Ag(QFS)1 target site. **(A)** A minimum of 50 U6::gRNA(*Dsx*-T1) DsRed+ males were crossed to a minimum of 50 *vas2::Cas9* YFP+ females. **(B)** The female offspring of the cross were examined for intersex mosaicism at the pupal and adult stage, visible by the development of male-like physical characteristics, depending on their genotype. The +/- signs indicate the presence (+) or absence (-) of the transgenes associated with each given fluorescent marker (DsRed/YFP).

**Supplementary Figure 6.**
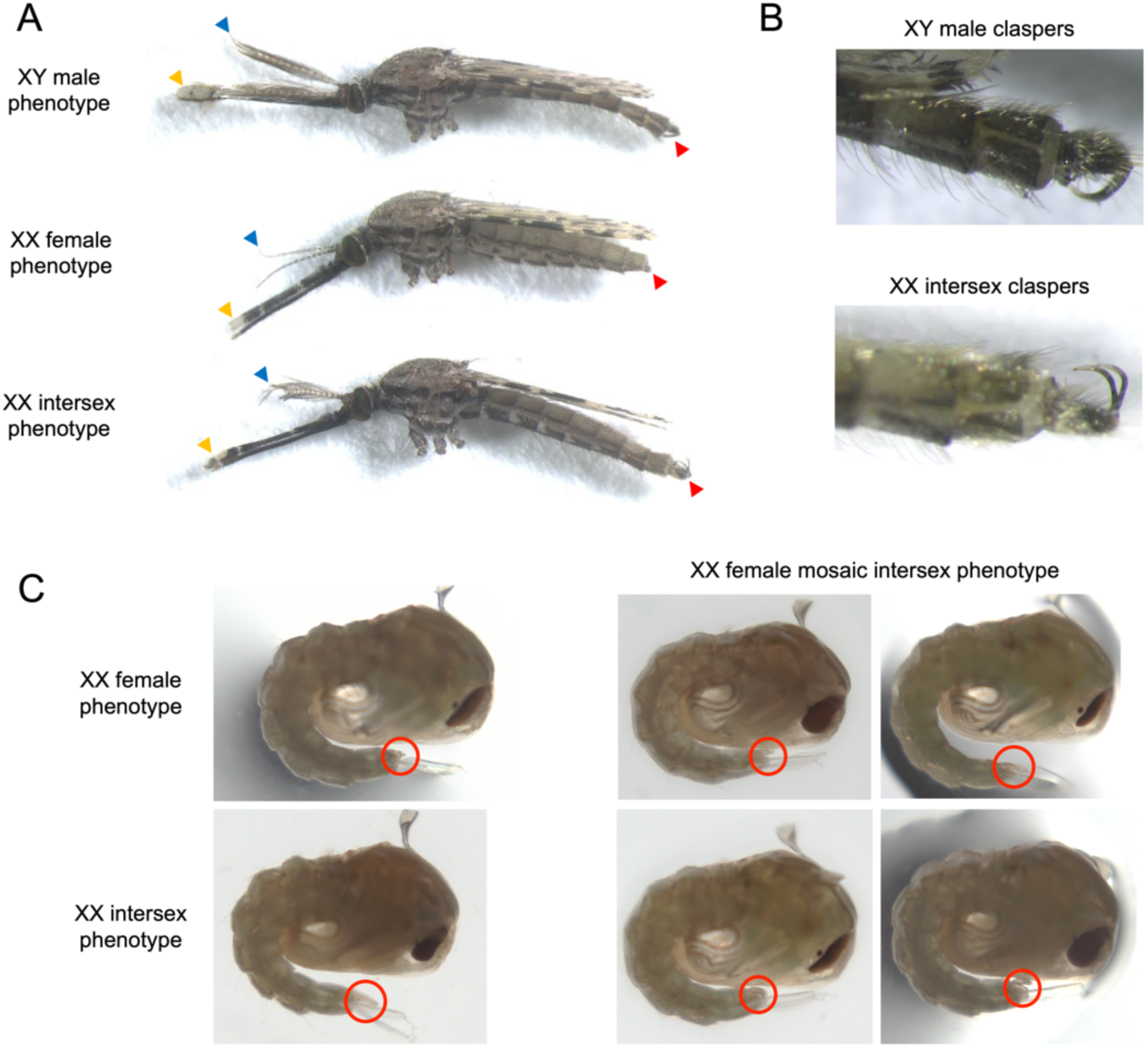
The intersex phenotype. **(A)** Intersex females develop semi-plumose antennae (blue arrows), resembling that of males; male-like palps, and a proboscis that is unable to draw blood (yellow arrows); and under-developed claspers, which females completely lack, facing upwards, instead of downwards like in males (red arrows). **(B)** Claspers rotate to face downwards in mature males, whereas intersex claspers remain facing upright. **(C)** Pupal genitalia are denoted using red circles. Fully intersex females develop male-like genitalia (uniform phenotype). Mosaic intersex individuals can be distinguished by their under-developed male-like genitalia (variable phenotype, showing different degrees of penetrance depending on the level of mosaicism).

**Supplementary Figure 7.**
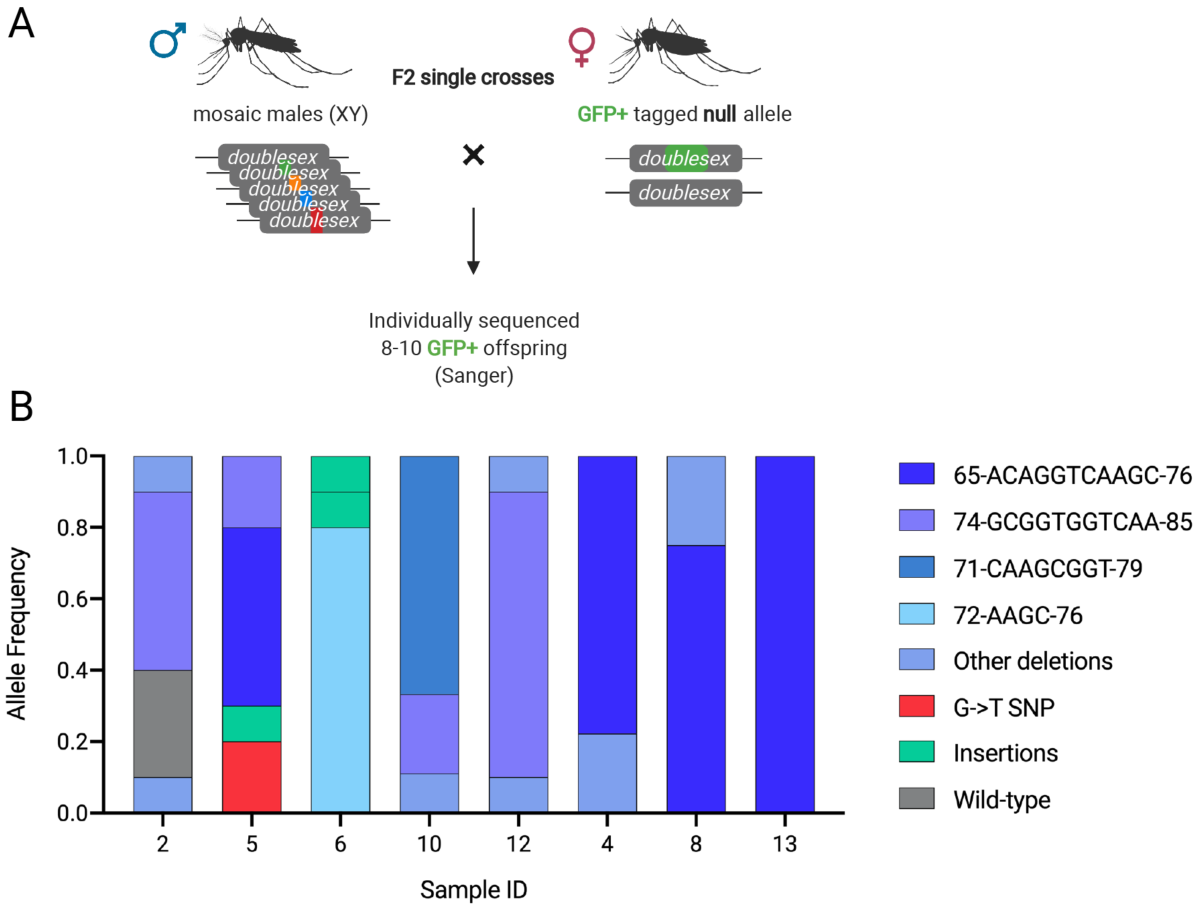
Distinct heritable mutations that occur at the Ag(QFS)1 target site through embryonic deposition of Cas9. **(A)** Schematic of F2 single crosses that were set up to answer how many distinct mutations each mutated male parent contributed to its offspring. 8 Mosaic males carrying mutations at *dsx*-T1, due to *vas*-Cas9 maternal deposition, were crossed to females carrying a GFP-tagged *dsxF* null allele, in single crosses. **(B)** The portion of distinct mutations inherited amongst their offspring, as determined by Sanger sequencing is shown (each cross is denoted with a distinct sample ID). 8-10 individuals were examined per clutch. In total,12 distinct mutant alleles were discovered in the 76 individual offspring that were sequenced collectively from all single crosses.

**Supplementary Figure 8.**
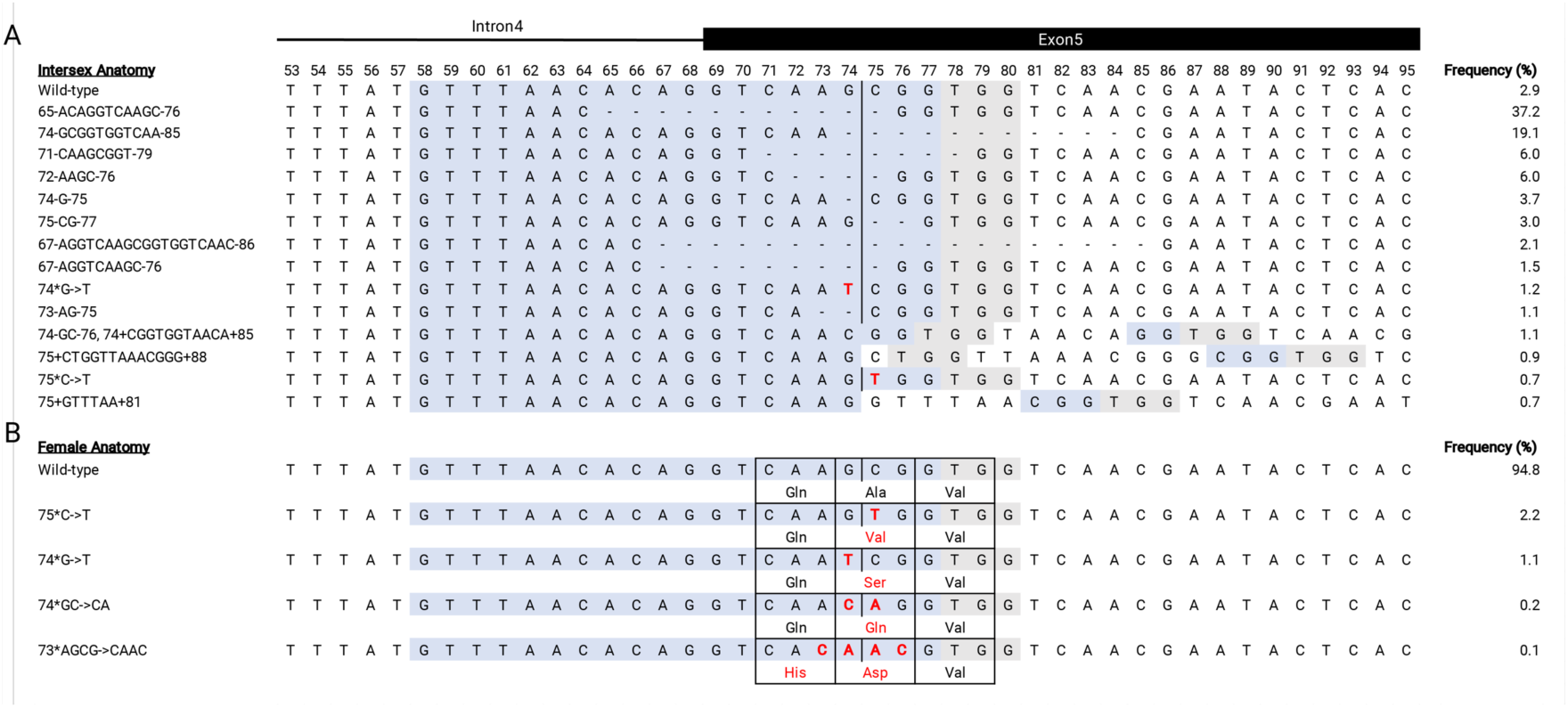
Sequence alignment of the alleles discovered through the mutagenesis screen at the Ag(QFS)1 target site. Deletions are denoted by dashes, and SNPs are depicted in red. The T1 gRNA spacer binding site is highlighted in blue and the PAM in grey. The frequency of each depicted allele is shown to the right. **(A)** Alleles discovered in anatomically intersex (sterile) GFP+ females. **(B)** Alleles discovered in GFP+ females that developed normally, as typical females. The amino acids encoded by codons carrying SNPs are also shown.

**Supplementary Figure 9.**
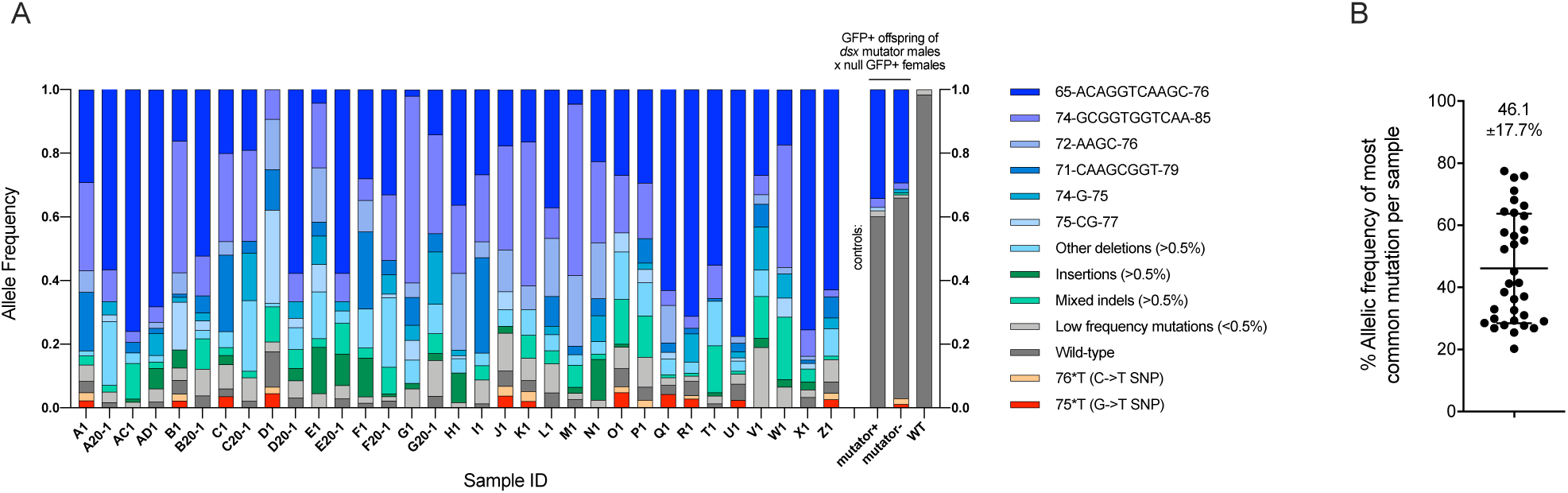
The types of Cas9-induced EJ mutations at the Ag(QFS)1 target site, recovered in the intersex fraction of F3 offspring from from distinct F2 crosses of the mutagenesis screen. **(A)** A minimum of 50 mosaic males containing a multitude of Cas9-induced EJ mutations were crossed to a minimum of 50 *dsx* null (*dsxF-*)-carrying females (GFP+)^7^. This cross was performed in replicate cages 33 times (cages A, B, C, D, E, F, G, H, I, J, K, L, M, N, O, P, Q, R, T, U, V,W, X, Z, AC, AD, A20, B20, C20, D20, E20, F20, G20). For each cross, 100 GFP+ intersex offspring were analysed through pooled amplicon Illumina sequencing. The bars show the relative portion of each mutation recovered in intersex individuals (A). The graph shows the allelic frequency of the most common mutation present in each intersex offspring pool (B). As controls, three pools of 100 non-sex-separated individuals were subjected to pooled amplicon Illumina sequencing. The first two pools contained the GFP+ offspring of a cross of 50 *dsx* mutator males (expressing both a *zpg-* Cas9 and a gRNA against the target site of Ag(QFS)1) to 50 null GFP+ females, in the absence of the maternal *vas2::Cas9* strain. Offspring that had also inherited the *dsx* mutator allele (mutator+), in addition to the *dsxF-* GFP+ null allele, were analysed separately from those that did not inherit the *dsx* mutator allele (mutator-). The third pool only contained wild-type individuals (WT).

**Supplementary Figure 10.**
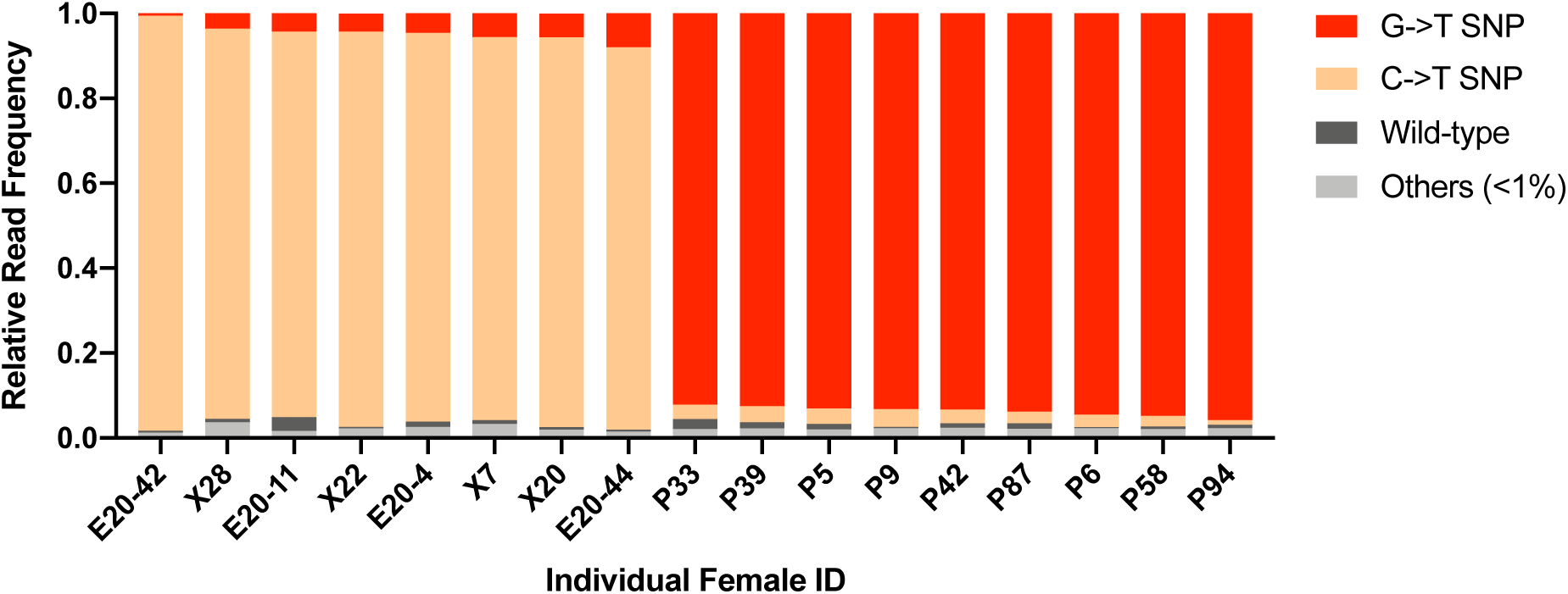
Amplicon sequencing of anatomical females carrying putative drive-resistant mutations. These samples were previously shown to carry a single allele paired to the null *dsx* mutation through Sanger sequencing: the C→T SNP (majority beige), the G→T SNP (majority red) or a WT allele (majority dark grey). F3 GFP+ individuals were batch-collected from large cages containing >500 mosquitoes and separated on a CO_2_ pad into three groups comprising of: (1) males, (2) anatomical females, and (3) anatomical intersex, for long-term storage (>6-12 months due to Covid-19 interruptions). Prior to gDNA extraction anatomical females were individually separated, however a low level of cross-contamination of the samples is possible.

**Supplementary Figure 11.**
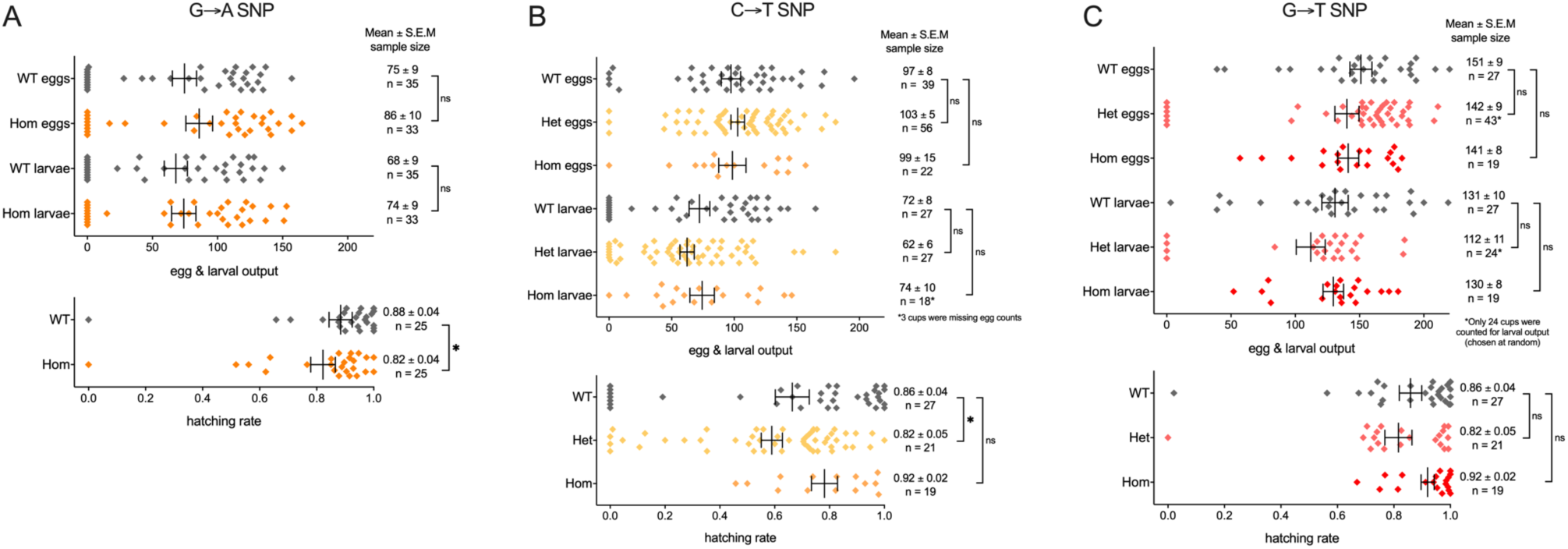
Fertility of females carrying each of the SNP variants engineered at the target site of Ag(QFS)1. **(A)** Fertility of females carrying the natural GèA SNP. The datasets did not pass the D’Agostino-Pearson normality test and therefore each experimental dataset was compared to the wild-type control using the Mann-Whitney non-parametric test. Ns = not significant (WT vs Hom egg output with p=0.3051, WT vs Hom larval output with p=0.6459), ∗ = significant with p=0.0463. **(B)** Fertility of females carrying the Cas9-induced CèT SNP. Only egg output datasets passed the D’Agostino-Pearson normality test and were analysed using ordinary ANOVA, allowing for multiple comparisons to the wild-type control. Larval output and hatching rate datasets were compared to the appropriate wild-type control using the Kruskall-Wallis non-parametric test. Ns = not significant (WT vs Het egg output with p=0.7869, WT vs Hom egg output with p=0.9946, WT vs Het larval output with p=0.4326, WT vs Hom larval output with p>0.9999, WT vs Hom hatching rate p>0.9999), ∗ = significant with p=0.0445. **(C)** Fertility of females carrying the Cas9-induced GèT SNP. The datasets did not pass the D’Agostino-Pearson normality test and therefore each experimental dataset was compared to the wild-type control using the Kruskall-Wallis non-parametric test, allowing for multiple comparisons. Ns = not significant (WT vs Het egg output with p>0.9999, WT vs Hom egg output with p>0.9999, WT vs Het larval output with p=0.4874, WT vs Hom larval output with p>0.9999, WT vs Het hatching rate with p=0.4863, WT vs Hom hatching rate with p=0.6497). Abbreviations: WT = wild-type, Het = heterozygous for each variant, Hom = homozygous for each variant.

**Supplementary Figure 12.**
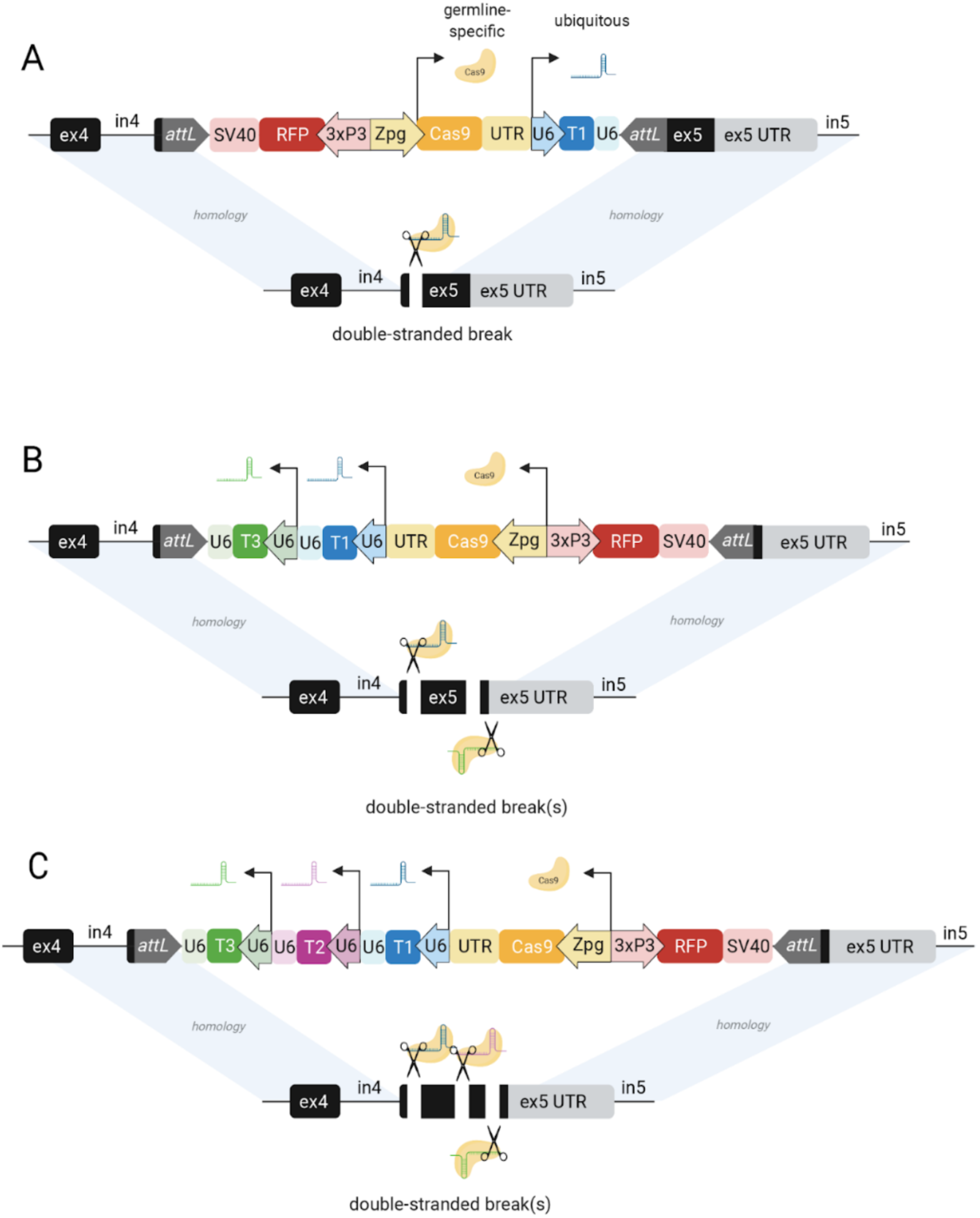
Schematics of the gene drive constructs targeting the *dsx* gene, tested in the present study. **(A)** The Ag(QFS)1 gene drive construct integrated in the same orientation as the *doublesex* gene. **(B)** The Ag(QFS)2 gene drive construct integrated in the reverse orientation with respect to *dsx*. **(C)** The Ag(QFS)3 gene drive construct integrated in the reverse orientation with respect to *dsx*. Gene drive components: att: = RMCE ruminant attachment sites, SV40 = viral terminator, RFP = DsRed fluorescent marker, 3xP3 = neuronal promoter, zpg = *zero population growth* promoter and untranslated region (UTR), Cas9 = human codon-optimised *Streptococcus pyogenes* Cas9 (SpCas9) gene, U6 = pol III promoter and terminator, T1/2/3 = gRNAs containing spacer sequences complementary to sites T1, T2 or T3.

**Supplementary Figure 13.**
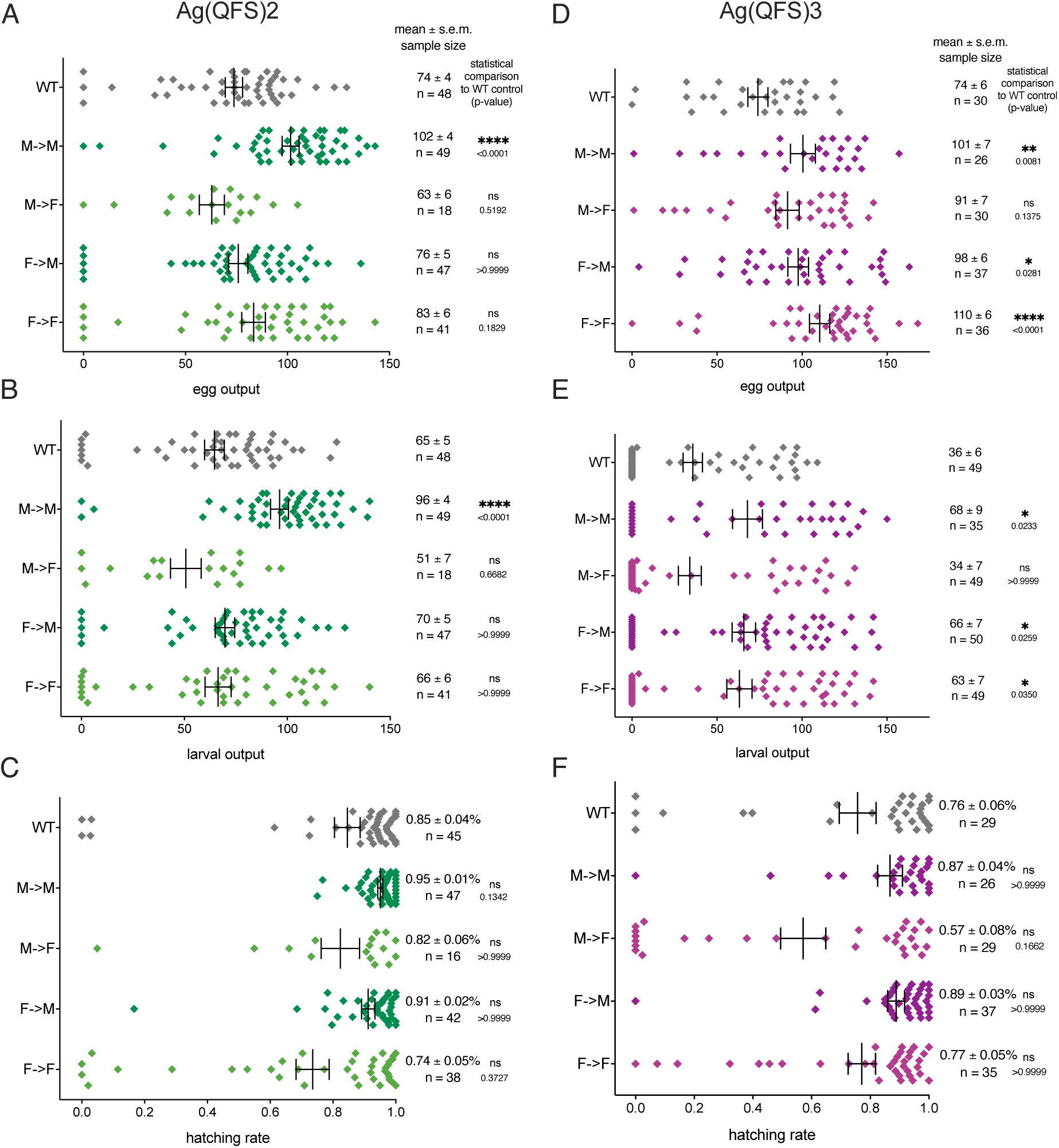
Fitness of Ag(QFS)2 (A-C) and Ag(QFS)3 (D-F) multiplexed gene drive carriers compared to wild-type controls. Male (M) and female (F) heterozygous gene drive carriers that inherited the paternally (M→M, M→F) or maternally (F→M, F→F) were crossed to wild-type, and the number of eggs (A, D) and larvae (B, E) produced per female parent were scored. The hatching rate of the eggs is also shown (C, F). Mean values, the standard error around the mean (S.E.M.), together with the sample size (n) and p-values derived from Kruskall-Wallis statistical comparisons to the wild-type control are shown to the right of each graph. Both mated and unmated females are included in this analysis.

**Supplementary Figure 14.**
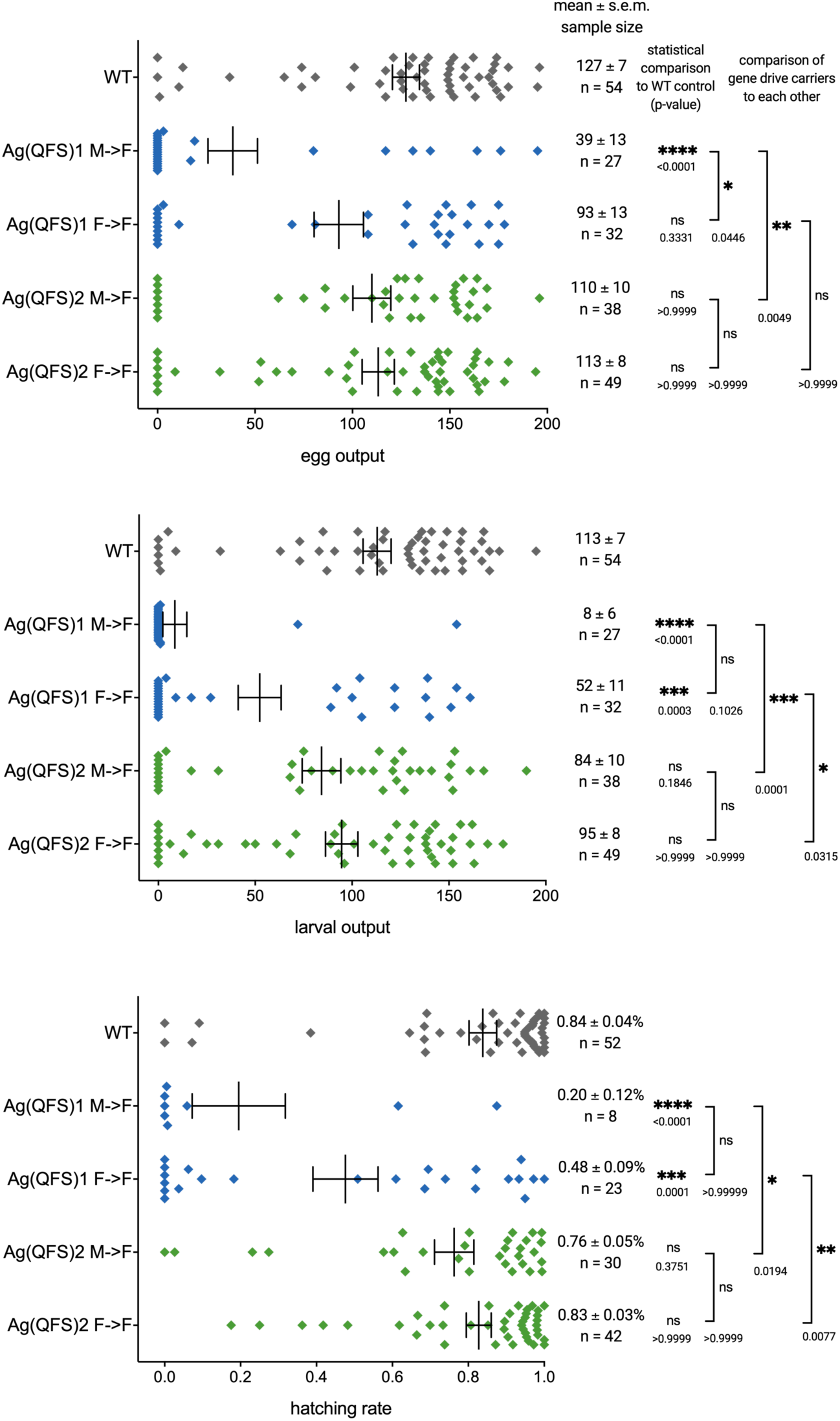
Fitness of Ag(QFS)2 gene drive carriers, compared to Ag(QFS)1. Female (F) heterozygous individuals carrying either Ag(QFS)1 or Ag(QFS)2, that had inherited each gene drive paternally (M→F) or maternally (F→F) were crossed to wild-type (WT), and the number of eggs (top) and larvae (middle) produced per female parent were scored. At the same time the eggs and larvae produced by a wild-type cross were scored to function as a control. The hatching rate of eggs is also shown (bottom). Mean values, the standard error around the mean (S.E.M.), sample size (n) and p-values derived from Kruskal-Wallis statistical comparisons to the WT control are shown to the right of each graph, as well as pairwise comparisons between specific groups as indicated by the brackets. Both mated and unmated females are included in this analysis.

**Supplementary Figure 15.**
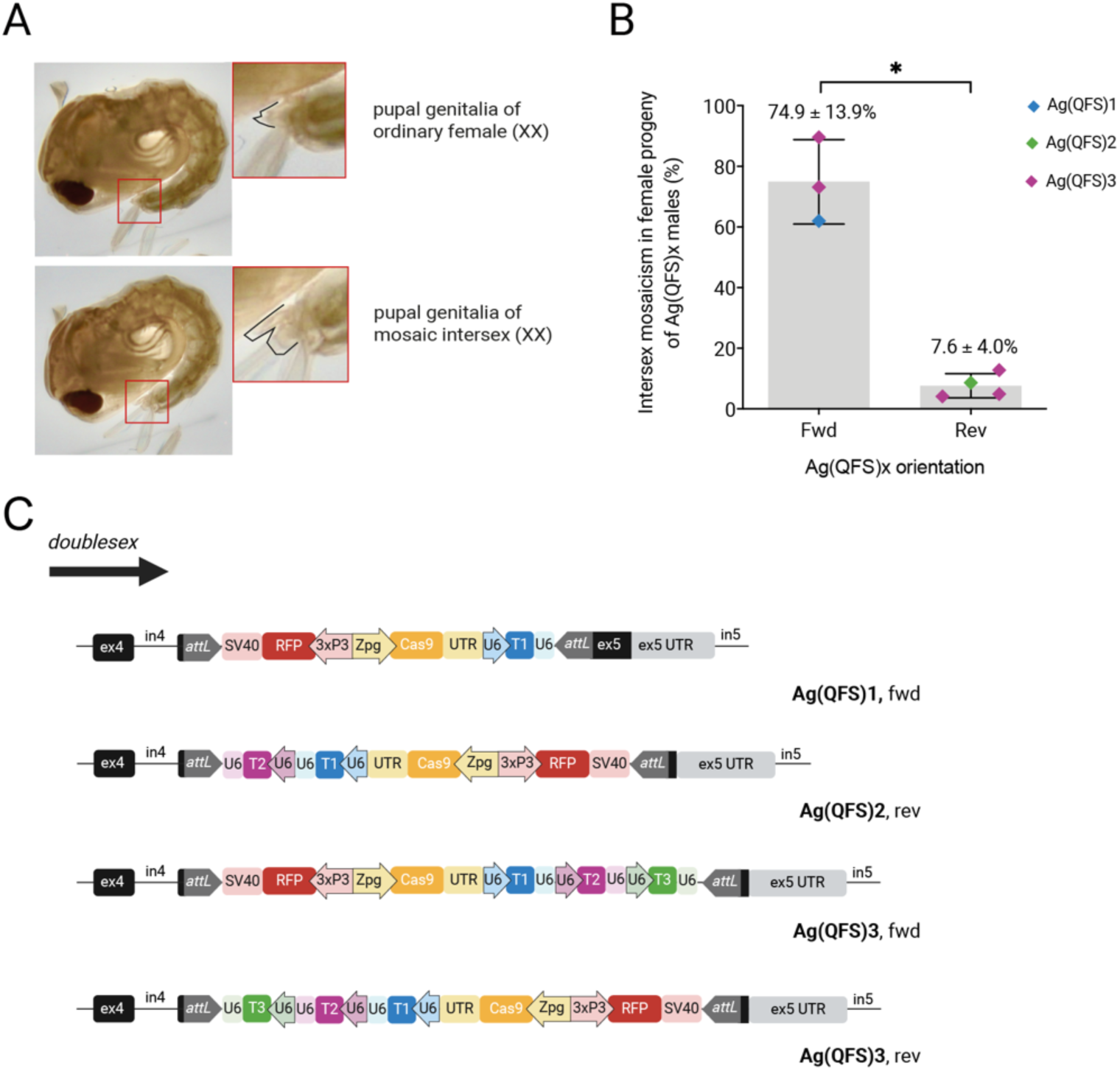
Intersex mosaicism in female QFS gene drive carriers. **(A)** Example of the female genitalia at the pupal stage, of phenotypic females (top) versus mosaic intersex females (bottom). **(B)** Percentage of intersex mosaics in the female offspring of male gene drive carriers of Ag(QFS)1, Ag(QFS)2 or Ag(QFS)3 that harboured the gene drive in the same (fwd) or reverse (rev) orientation in the genome, with respect to *dsx.* Means and standard deviations are shown above the graph and indicated by the error bars. A t-test with Welch’s correction to account for unequal standard deviation in the two datasets was performed with p-value = 0.0104. Note that each data point originated from an independently generated strain (i.e. isolated from a different founder). **(C)** Illustrations of the gene drive integration mode with respect to *dsx* (fwd or rev) for each of the gene drives examined.

**Supplementary Figure 16.**
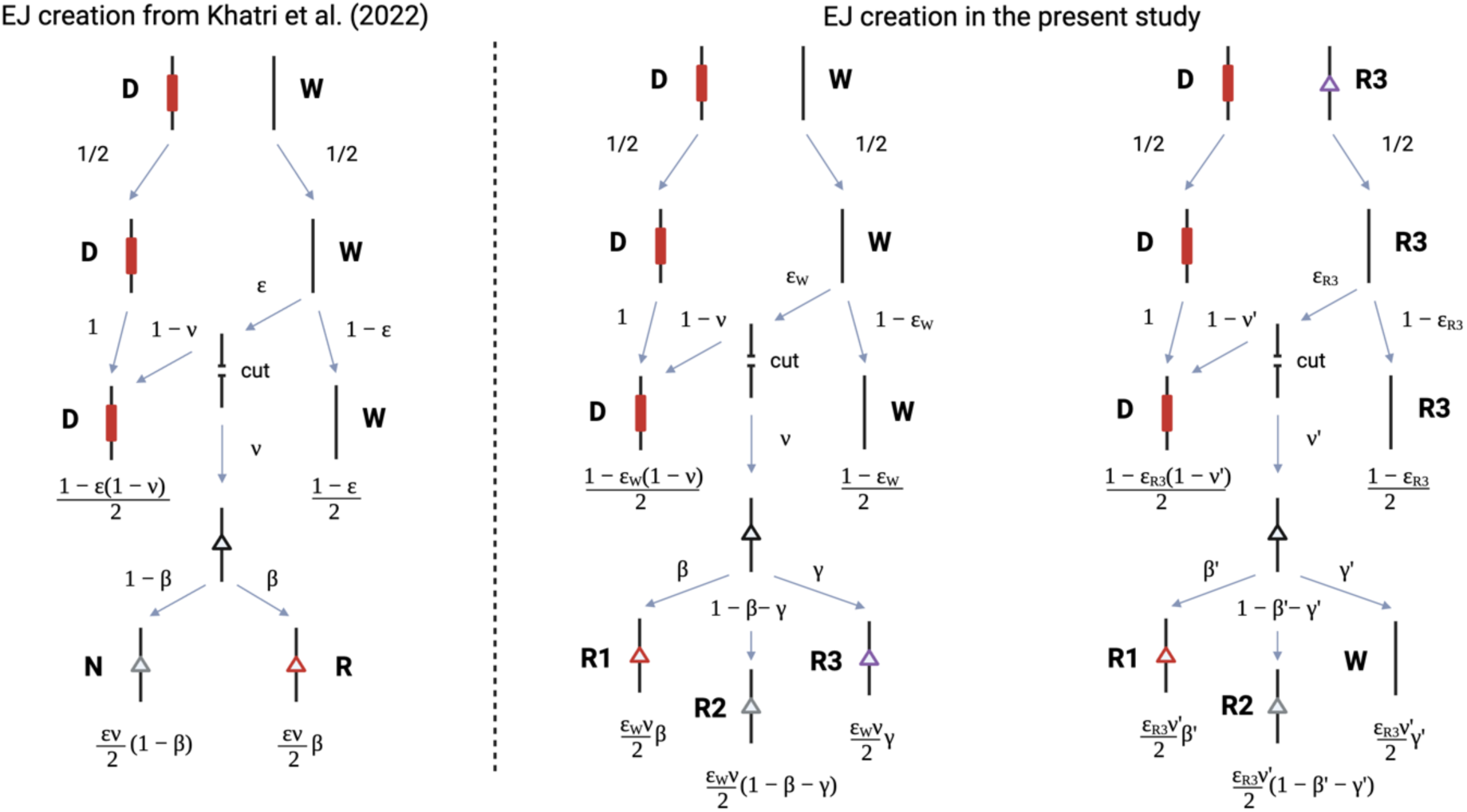
Diagram of resistant allele creation during gametogenesis, as incorporated into our modelling framework. Abbreviations: EJ = end-joining, D = drive allele,W= wild-type allele, N = non-functional EJ mutation, R = functional resistant mutation, R1 = functional resistant mutation, R2 = non-functional resistant mutation, R3 = functional partially resistant mutation. Model parameters: ε = cleavage efficiency, ε_W_ = cleavage efficiency of wild-type, ε_R3_ = cleavage efficiency of R3, ν = fraction of mutated alleles, β = fraction that are R1, γ = fraction that are R3, β’ = fraction that are R1, γ’ = fraction that are wild-type.

**Supplementary Table 1.** The number of genetic females assessed as part of the mutagenesis screen, and the portion that carried putative resistant mutations.

| GFP+ genetic females collected |  | N | % |
| --- | --- | --- | --- |
| Total |  | 10,083 | 100.0 |
| Female anatomy |  | 1,031 | 10.2 |
| Intersex anatomy |  | 9,052 | 89.8 |
| Sanger-sequenced anatomical females |  | N | % |
| Total |  | 852 | 100.0 |
| Wild-type | Blood-fed | 629 | 94.8 |
|  | Non blood-fed | 107 |  |
|  | Unchallenged | 72 |  |
| C->T SNP | Blood-fed | 14 | 2.2 |
|  | Non blood-fed | 5 |  |
| G->T SNP | Blood-fed | 9 | 1.1 |
|  | Non blood-fed | 0 |  |
| Others modified | Blood-fed | 2 | 0.4 |
|  | Non blood-fed | 1 |  |
| Mixed modified alleles* | Blood-fed | 10 | 1.5 |
|  | Non blood-fed | 3 |  |

\*Likely resulting from mosaicism due to residual *zpg*-expressed Cas9 activity.

**Supplementary Table 2.** Modelling parameters. Also refer to Supplementary Figure 16 to understand how the modelling parameters relate to each other.

| Parameters | Description | Values |
| --- | --- | --- |
| $\varepsilon_W$ | Cleavage rate of WT that leads to non-WT allele | 0.966 |
| $1 - \varepsilon_W$ | Frequency of unmodified WT allele after exposure to nuclease | 0.034 |
| $\nu$ | Frequency of WT allele conversion to EJ mutant allele | 0.035 |
| $1 - \nu$ | Fraction of drive alleles amongst those cleaved | 0.965 |
| $\beta$ | Fraction of EJ mutations that become an R1 allele | 0.0016 |
| $1 - \beta - \gamma$ | Fraction of EJ mutations that become an R2 allele | 0.996 |
| $\gamma$ | Fraction of EJ mutations that become an R3 allele | 0.0025 |
| $\varepsilon_{R3}$ | Cleavage rate of R3 alleles | 0.416 |
| $1 - \varepsilon_{R3}$ | Frequency of unmodified R3 alleles after exposure to nuclease | 0.584 |
| $\nu'$ | Frequency of R3 conversion to EJ mutant allele | 0.000 |
| $1 - \nu'$ | Fraction of drive alleles amongst those cleaved | 1.000 |
| $\beta'$ | Fraction of EJ mutations that become an R1 allele | 0.000 |
| $1 - \beta' - \gamma'$ | Fraction of EJ mutations that become an R2 allele | 1.000 |
| $\gamma'$ | Fraction of EJ mutations that become a WT allele | 0.000 |
| - | Fraction of non-drive, cleaved WT alleles that are converted to EJ mutant alleles | 0.500 |
| - | Fraction of non-drive, cleaved R3 alleles that are converted to EJ mutant alleles | 0.020 |

